# The Ski2 helicase ASCC3 unwinds DNA upon fork stalling to control replication stress responses

**DOI:** 10.1101/2025.07.24.666583

**Authors:** Shixin Cui, Nicole L. Batenburg, Yan Coulombe, Aruna Arumugam, John R. Walker, Sadaf Valeh Sheida, Anja-Katrin Bielinsky, Markus C. Wahl, Jean-Yves Masson, Xu-Dong Zhu

## Abstract

The activating signal co-integrator 1 complex subunit 3 (ASCC3), the largest subunit of ASCC, is one of two Ski2-like helicases with duplicated helicase cassettes encoded by the human genome. ASCC3 has been implicated in transcriptional regulation, alkylation damage repair, and ribosome quality control. In addition, published proteomics studies suggest that ASCC3 is associated with stalled forks. However, little is known about its role in replication stress. Here we report that ASCC3 is recruited to stalled forks by its binding partner ASCC2, whose recruitment to stalled forks is dependent upon both its ubiquitin binding activity and polyubiquitylation of PCNA at K164 that is catalyzed by SHPRH, HLTF, and RFWD3. In response to replication stress. ASCC3 unwinds DNA, generating both parental and nascent ssDNA. This unwinding activity is required to promote fork reversal. We demonstrate that ASCC3 also interacts with RPA and stimulates RPA accumulation on ssDNA upon replication stress, promoting ATR activation. Furthermore, ASCC3 functions to antagonize RAD51-mediated recombination and prevents accumulation of chromosome breaks/gaps and mis-segregation in response to replication stress. Our work underscores a previously uncharacterized but critical role of ASCC3 in controlling multiple replication stress responses to maintain genomic stability.

## INTRODUCTION

When encountering genotoxins, progression of DNA replication forks is slowed or stalled^1,2^, a situation generally referred to as replication stress. Replication stress represents a major source of genomic instability that can drive tumorigenesis and yet at the same time is exploited by many clinically relevant chemotherapeutics^2–4^. Human cells have evolved several interconnected responses to replication stress^5–7^. Upon fork stalling, the uncoupling of leading and lagging strand DNA synthesis generates single-stranded DNA (ssDNA). Persistent binding of the ssDNA by replication protein A (RPA) activates ATR serine/threonine kinase (ATR), which suppresses the firing of dormant origins and induces cell cycle arrest^5,8^. In addition, stalled forks can be remodeled into reversed forks, a process known as fork reversal^5,6,9^, which slows down fork progression to enable lesion repair. Furthermore, several DNA damage tolerance (DDT) pathways are activated^4,5^, including translesion DNA synthesis (TLS), template switching, and fork repriming that uses a specialized enzyme called primase and DNA-directed polymerase (PRIMPOL). PRIMPOL skips DNA lesions to re-initiate DNA synthesis, leaving a ssDNA gap that is repaired post-replicatively^10–12^. Fork reversal is thought to inhibit PRIMPOL-mediated fork repriming^13,14^. A precursor of fork reversal is the formation of ssDNA upon fork stalling^15–17^. Several RecQ helicase family members have been implicated in fork reversal^18,19^, including BLM which has been directly implicated in fork reversal in cells^20,21^. The UvrD family member F-box DNA helicase 1 (FBH1) is also required for fork reversal^22^. However, whether other helicase families might unwind DNA upon replication stress to facilitate fork reversal remains largely elusive.

Ubiquitylation of proliferating cell nuclear antigen (PCNA), a homotrimer that functions as a sliding clamp for DNA polymerases during DNA synthesis and repair, represents a major signaling event in response to replication stress^23^. PCNA K164 is monoubiquitylated by the E3 ligase RAD18^24^ and this monoubiquitylation directs DDT pathway choice to translesion DNA synthesis (TLS). PCNA K164 can also be modified by K63-linked polyubiquitin chains. Several E3 ligases, such as SNF2 histone linker PHD ring helicase (SHPRH), helicase like transcription factor (HLTF), and ring finger and WD repeat domain 3 (RFWD3), catalyze this polyubiquitylation^25–27^, channeling DDT pathway choice to template switching. Aside from DDT pathway choice, PCNA ubiquitylation at K164 has also been implicated in regulating fork reversal^23,28,29^.

Activating signal co-integrator 1 complex (ASCC), which consists of ASCC1, ASCC2, ASCC3, and TRIP4 subunits^30–32^, was first reported to function as a transcriptional coactivator of nuclear receptors^30^. Subsequently, ASCC has been implicated in alkylation damage repair^31,32^, ribosome quality control^33,34^, and RNA splicing^35^. Several published studies provide compelling evidence that ASCC may be involved in the regulation of DNA replication. Firstly, ASCC3 is present in a list of proteins whose depletion is thought to enhance DNA re-replication^36^. Secondly, ASCC subunits are detected at both normal and stalled/broken DNA forks^37–39^. Thirdly, ASCC subunits are found as part of the PCNA interactome during unperturbed and damage-induced conditions^40^. However, the function that ASCC may carry out at stalled forks remains uncharacterized.

ASCC2 and ASCC3 are considered as core subunits of the ASCC complex since loss of either ASCC2 or ASCC3 leads to destabilization of ASCC whereas loss of either ASCC1 or TRIP4 does not^33^. ASCC2 contains a CUE (coupling of ubiquitin conjugation to ER degradation) domain, which binds K63-linked ubiquitin chains^32^. ASCC3, which interacts with all three other ASCC subunits^32,41–43^, is one of the two enzymes encoded by the human genome that contain duplicated Ski2-like helicase cassettes arranged in tandem^42^. The other is the RNA helicase bad response to refrigeration 2 homolog (BRR2). ASCC3 is a 3’-5’ DNA helicase that only unwinds double-stranded DNA (dsDNA) containing a 3’-overhang in vitro and it shows no helicase activity towards dsDNA with a 5’-overhang or blunt ends^31^. It has been suggested that ASCC3 is recruited by ASCC2, which senses E3 ubiquitin ligase RNF113A-mediated ubiquitin signaling, to DNA alkylation damage sites in G1, generating ssDNA substrates for the alkylation repair enzyme AlkB homolog 3, alpha-ketoglutarate dependent dioxygenase (ALKBH3)^32^. However, whether ASCC3 plays a role in unwinding DNA in S phase in response to replication stress is unknown.

In this report, we have discovered that ASCC3 is recruited to stalled forks by ASCC2, whose recruitment to stalled forks requires both its ubiquitin binding activity and polyubiquitylation of PCNA at K164. Our finding suggests that ASCC3 catalyzes DNA unwinding upon replication stress and that this unwinding activity is a prerequisite for fork reversal. In addition, ASCC3 interacts with RPA and stimulates RPA accumulation on ssDNA, thereby promoting efficient ATR activation. Furthermore, ASCC3 functions to antagonize RAD51-dependent recombination and prevents accumulation of chromosome breaks/gaps and mis-segregation. Our finding underscores ASCC3 as a critical regulator of genomic stability by controlling multiple replication stress responses.

## RESULTS

### ASCC2 and ASCC3 are recruited to stalled forks

ASCC2 and ASCC3 are detected at normal and stalled/broken DNA forks in several published proteomic studies^37–39^. To investigate whether ASCC plays a role in replication stress, we knocked out, via CRISPR-Cas9, ASCC2 or ASCC3 in U2OS cells (Fig. 1A). We observed that loss of ASCC2 impaired the level of ASCC3 and vice versa (Fig. 1A), in agreement with a published finding^33^. To confirm whether ASCC2 and ASCC3 are recruited to stalled forks, we turned to a proximity ligation assay (PLA) that allows one to measure *in situ* interactions of protein-protein or protein-DNA interactions. We first pulse-labeled U2OS parental (WT) and ASCC2-KO cells with EdU for 10 min prior to HU treatment. Treatment with HU led to an increase in the formation of PLA foci between endogenous ASCC2 and EdU (Fig. 1B and 1C). This increase was abolished by loss of ASCC2 (Fig. 1B and 1C). To address a concern that a change in EdU incorporation prior to treatment with HU might affect EdU-ASCC2 PLA foci formation, we measured the formation of EdU-EdU PLA foci in both U2OS WT and ASCC2-KO cells. ASCC2 loss had no effect on EdU incorporation (Supplementary Fig. S1A and S1B), suggesting that it is unlikely that a change in EdU incorporation is responsible for the observed loss of EdU-ASCC2 PLA foci formation in U2OS ASCC2-KO cells. To address another concern that the observed EdU-ASCC2 PLA foci formation might be due to non-specific effects associated with PLA assays, we performed several control experiments, including no EdU incorporation, no primary antibodies, a primary antibody against a cytoplasmic protein such as fatty acid synthase (FASN), as well as a primary antibody against a protein known to be associated with stalled forks such as PCNA. We found that omitting EdU or primary antibodies (1° Abs) led to little detectable PLA foci formation (Fig. 1D and Supplementary Fig. S1C). FASN also failed to form PLA foci with EdU in HU-treated U2OS cells (Fig. 1D and Supplementary Fig. S1C). These results altogether further demonstrate that the observed EdU-ASCC2 PLA foci formation is unlikely due to non-specific effects. We observed that the number of EdU-ASCC2 PLA foci in HU-treated U2OS cells was much less than that of EdU-PCNA PLA foci (Fig. 1D and Supplementary Fig. S1C), suggesting that ASCC2 is recruited to stalled forks either at a low level or in a transient manner.

**Figure 1.**
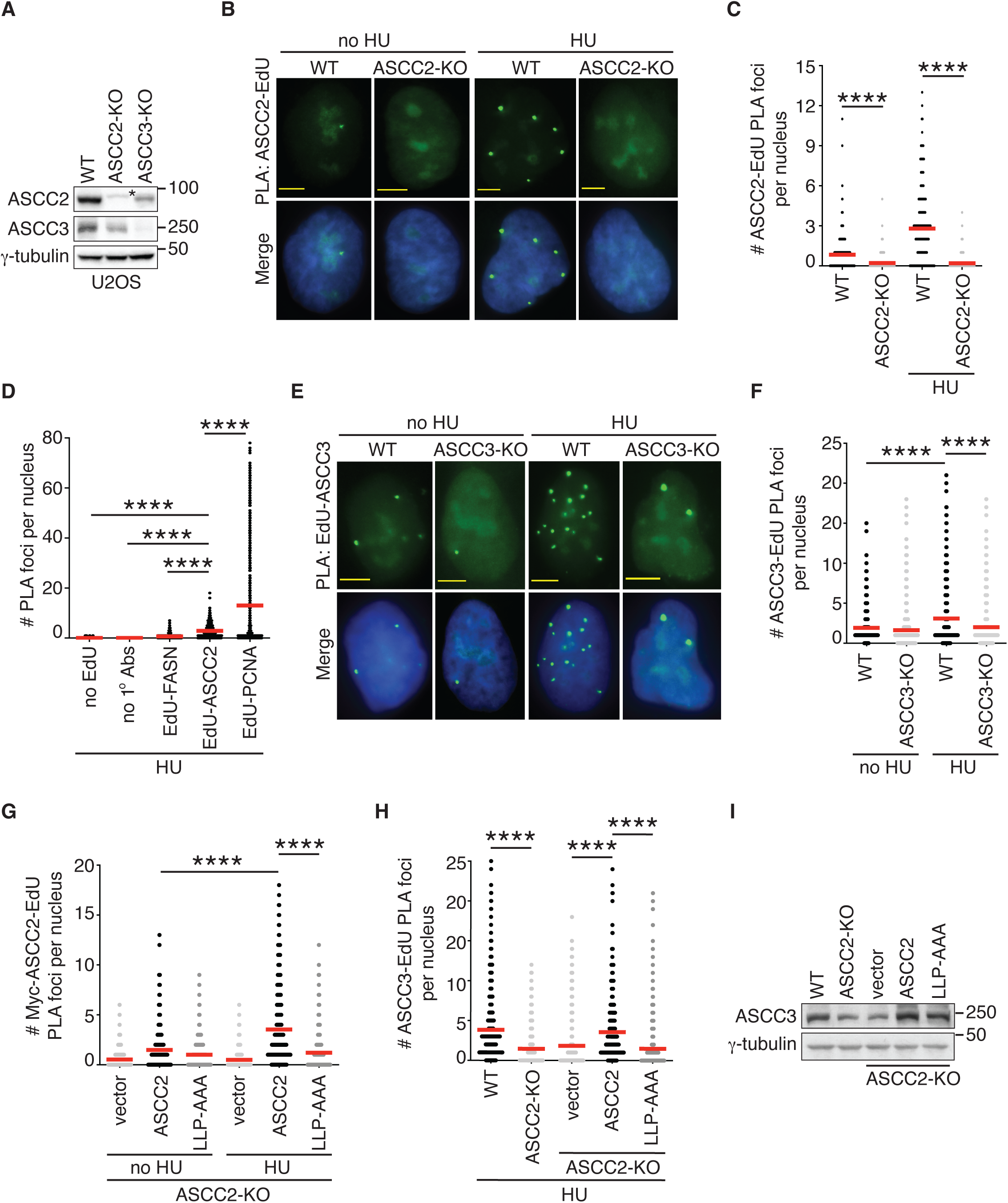
ASCC2 recruits ASCC3 to stalled forks in a manner dependent upon ASCC2’s ubiquitin binding activity. (**A**) Western blot analyses of U2OS WT, ASCC2-KO, and ASCC3-KO cells. Immunoblotting was performed with anti-ASCC2, anti-ASCC3, and anti-γ-tubulin antibodies. The γ-tubulin blot was used as a loading control in this and subsequent figures except for where the α-tubulin blot was used as a loading control. The asterisk (*) indicates a non-specific band. (**B**) Representative images of endogenous ASCC2-EdU PLA foci formation in U2OS WT and ASCC2-KO that were either treated without or with 4 mM HU for 4 h. Nuclei were stained with DAPI in blue in this and subsequent figures. Scale bars in this and subsequent figures: 5 μm. (**C**) Quantification of endogenous ASCC2-EdU PLA foci formation from (B). The PLA experiments were performed independently twice with reproducible data in this, 1F, 1G, and 1H panels. Data from one representative experiment are shown as scatter plot graphs with the mean indicated in this and subsequent panels. A total of 492-506 cells per condition were analyzed. The *P*-value was determined using a non-parametric Mann-Whitney rank-sum *t*-test in this and subsequent panels. \*\*\*\**P*<0.0001. (**D**) Quantification of PLA foci formation in U2OS treated with 4 mM HU for 4 h. PLA assays were performed in several conditions as indicated. This PLA experiment was performed once. A total of 492-515 cells per condition were analyzed. \*\*\*\**P*<0.0001. (**E**) Representative images of endogenous ASCC3-EdU PLA foci formation in no HU- or HU-treated U2OS WT and ASCC3-KO cells. (**F**) Quantification of endogenous ASCC3-EdU PLA foci formation from (E). A total of 1389-1396 cells per condition were analyzed. \*\*\*\**P*<0.0001. (**G**) Quantification of Myc-tagged ASCC2-EdU PLA foci formation in no HU- or HU-treated U2OS ASCC2-KO cells expressing various Myc-ASCC2 alleles as indicated. A total of 603-617 cells per condition were analyzed. \*\*\*\**P*<0.0001. (**H**) Quantification of endogenous ASCC3-EdU PLA foci formation in no HU- or HU-treated U2OS ASCC2-KO cells stably expressing various Myc-ASCC2 alleles as indicated. A total of 806-816 cells per condition were analyzed. \*\*\*\**P*<0.0001. (**I**) Western blot analyses of U2OS ASCC2-KO cells stably expressing various Myc-ASCC2 alleles as indicated. Immunoblotting was performed with anti-ASCC3 and anti-γ-tubulin antibodies.

We also investigated whether endogenous ASCC3 is recruited to stalled forks. We pulse-labeled U2OS WT and ASCC3-KO cells with EdU for 10 min prior to HU treatment. As with ASCC2, treatment with HU also led to an increase in the formation of PLA foci between endogenous ASCC3 and EdU (Fig. 1E and 1F), which was abrogated by ASCC3 loss (Fig. 1E and 1F). These results altogether suggest that both ASCC2 and ASCC3 are recruited to stalled forks, agreeing with published proteomic studies^37–39^.

### ASCC2 relies on its ubiquitin binding activity to recruit ASCC3 to stalled forks

ASCC2 contains a CUE domain that binds K63-linked ubiquitin chains^32^. It has been reported that triple L^478^L^479^P^480^-AAA mutations in the CUE domain of ASCC2 abolish its ubiquitin binding activity^32^. To investigate whether ASCC2 requires its ubiquitin binding activity to be recruited to stalled forks, we generated U2OS ASCC2-KO cells stably expressing the vector alone, Myc-tagged ASCC2 or Myc-tagged ASCC2 carrying L^478^L^479^P^480^-AAA mutations (Myc-ASCC2-LLP-AAA) (Supplementary Fig. S2A). Analysis of PLA assays revealed that while Myc-ASCC2 formed HU-induced PLA foci with EdU, Myc-ASCC2-LLP-AAA failed to do so (Fig. 1G and Supplementary Fig. S2B). The latter was unlikely due to a defect in protein expression since Myc-ASCC2-LLP-AAA was expressed comparably to Myc-ASCC2 (Supplementary Fig. S2A). These results altogether suggest that ASCC2 relies on its ubiquitin binding activity to be recruited to stalled forks.

As ASCC3 is a DNA helicase, we investigated whether ASCC3 requires its helicase activity to be recruited to stalled forks. We performed rescue experiments in U2OS ASCC3-KO cells transfected with the vector alone, Myc-tagged ASCC3 or Myc-tagged ASCC3 carrying a previously reported helicase dead G1354D mutation^31^ (Supplementary Fig. S2C). We found that the helicase-dead G1354D mutation did not affect the ability of Myc-ASCC3 to form HU-induced PLA foci with EdU (Supplementary Fig. S2D and S2E), suggesting that the helicase activity of ASCC3 is dispensable for its recruitment to stalled forks.

To investigate whether ASCC3 is recruited by ASCC2 to stalled forks, we measured HU-induced ASCC3-EdU PLA foci formation in ASCC2-KO cells. Loss of ASCC2 abolished the number of HU-induced ASCC3-EdU PLA foci (Fig. 1H). Formally, it was possible that this abolishment might be due to a loss of ASCC3 in ASCC2-KO cells since knockout of ASCC2 results in a reduction in the level of ASCC3 (Fig. 1A). To address this possibility, we leveraged the ubiquitin binding-dead Myc-ASCC2-LLP-AAA mutant since this mutant was able to restore the level of endogenous ASCC3 in ASCC2-KO cells (Fig. 1I). Despite this restored ASCC3 level, the number of HU-induced endogenous ASCC3-EdU PLA foci remained impaired in ASCC2-KO cells expressing Myc-ASCC2-LLP-AAA (Fig. 1H), suggesting that ASCC3 is recruited by ASCC2 to stalled forks and that this recruitment requires ASCC2’s ubiquitin binding activity.

### Ubiquitylation of PCNA at K164 mediates recruitment of ASCC2 to stalled forks

A well-characterized K63-linked ubiquitin signaling event at stalled forks is polyubiquitylation of PCNA at K164^27^. SHPRH, HLTF, and RFWD3 are three E3 ubiquitin ligases that are reported to polyubiquitylate PCNA at K164^25–27^. As ASCC2’s CUE domain preferentially binds K63-linked ubiquitin chains^32^, we investigated whether these three E3 ligases might mediate ASCC2 recruitment to stalled forks. We knocked them down individually in U2OS cells (Supplementary Fig. S3A-S3C). While each depletion did not affect the level of ASCC2 (Supplementary Fig. S3D), it abrogated the formation of HU-induced ASCC2-EdU PLA foci in U2OS cells (Fig. 2A, Supplementary Fig. S3E), suggesting that SHPRH, HLTF, and RFWD3 act coordinately to mediate recruitment of ASCC2 to stalled forks. To further investigate whether E3 ligase activity is required for mediating ASCC2 recruitment to stalled forks, we focused on SHPRH. Using CRISPR-Cas9 genomic editing, we knocked out SHPRH in U2OS cells (Fig. 2B). Like depletion of SHPRH, knockout of SHPRH abolished the formation of HU-induced ASCC2-EdU PLA foci (Fig. 2C and 2D) but it did not affect the level of ASCC2 nor ASCC3 (Fig. 2B). Rescue experiments revealed that overexpression of Myc-SHPRH in SHPRH-KO cells restored HU-induced ASCC2-EdU PLA foci formation whereas this restoration was not detected in SHPRH-KO cells overexpressing Myc-SHPRH carrying a previously reported ubiquitin ligase deficient C1432A mutation^44^ (Myc-SHPRH-C1432A) (Fig. 2E). Myc-SHPRH-C1432A was expressed at a comparable level to Myc-SHPRH (Supplementary Fig. S3F). These results suggest that SHPRH E3 ligase activity is required to recruit ASCC2 to stalled forks.

**Figure 2.**
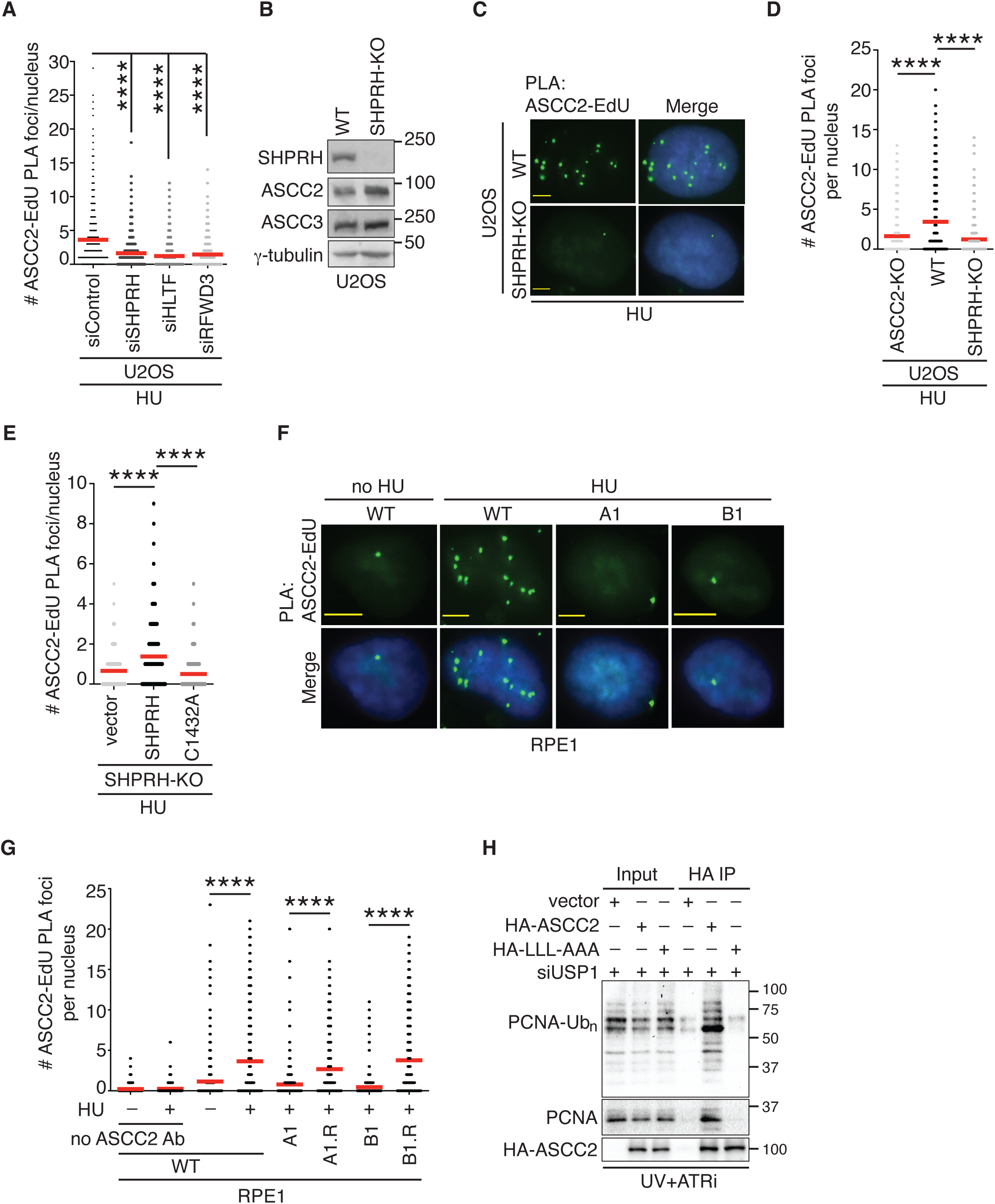
ASCC2 interacts with polyubiquitylated PCNA and is recruited by PCNA ubiquitylated at K164 to stalled forks. (**A**) Quantification of ASCC2-EdU PLA foci formation in HU-treated U2OS cells transfected with siControl or siRNA against SHPRH, HLTF or RFWD3. The PLA experiments were performed twice independently with reproducible data in this, 2D, and 2E panels. Data from one representative experiment are shown as scatter plot graphs with the mean indicated in this and subsequent panels. A total of 805-846 cells per condition were analyzed. The *P*-value was determined using a non-parametric Mann-Whitney rank-sum *t*-test in this and subsequent panels. \*\*\*\**P*<0.0001. (**B**) Western blot analyses of U2OS WT and SHPRH-KO cells. Immunoblotting was performed with anti-SHPRH, anti-ASCC2, anti-ASCC3, and anti-γ-tubulin antibodies. (**C**) Representative images of ASCC2-EdU PLA foci in HU-treated U2OS WT and SHPRH-KO cells. (**D**) Quantification of ASCC2-EdU PLA foci formation in HU-treated U2OS WT and SHPRH-KO cells from (C). U2OS ASCC2-KO cells were included as a control. A total of 817-822 cells per condition were analyzed. \*\*\*\**P*<0.0001. (**E**) Quantification of ASCC2-EdU PLA foci formation in HU-treated U2OS SHPRH-KO cells expressing various Myc-SHPRH alleles as indicated. A total of 827-832 cells per condition were analyzed. \*\*\*\**P*<0.001. (**F**) Representative images of ASCC2-EdU PLA foci in no HU- or HU-treated RPE1 parental (WT), A1 (PCNA-K164R), and B1 (PCNA-K164R) cells. (**G**) Quantification of ASCC2-EdU PLA foci formation in RPE1 WT, A1, B1, A1 revertant (A1.R, PCNA-K164K), and B1 revertant (B1.R, K164K) cells that were treated with or without HU. This PLA experiment was done once. A total of 525-540 cells per condition were analyzed. \*\*\*\**P*<0.0001. (**H**) Coimmunoprecipitation with anti-HA antibody in UV-irradiated, ATRi-treated and USP1-depleted HEK293T cells expressing the vector alone, HA-tagged ASCC2, or HA-tagged ASCC2 carrying LLL-AAA mutations. Immunoblotting was performed with anti-Ubiquityl-PCNA (K164), anti-PCNA, and anti-HA antibodies.

To investigate whether ubiquitylation of PCNA at K164 mediates recruitment of ASCC2 to stalled forks, we turned to two pairs of isogenic cell lines that have recently been reported^45^. The first pair are RPE1-A1 and RPE1-A1-R whereas the second pair are RPE-B1 and RPE1-B1-R. RPE1-A1 and RPE-B1 are two clones that each contain a knock-in PCNA K164R mutation whereas RPE1-A1-R and RPE-B1-R are revertant cell lines in which the K164R substitutions have respectively been reversed back to K164K^45^. Analysis of PLA assays revealed that like U2OS cells, RPE1 parental (WT) cells exhibited HU-induced ASCC2-EdU PLA foci formation (Fig. 2F and 2G), which was specific to ASCC2 since little ASCC2-EdU PLA foci formation was detected when the ASCC2 antibody was omitted in PLA assays (Fig. 2G). These results suggest that ASCC2 recruitment to stalled forks is not specific to U2OS cells. The knock-in PCNA K164R mutation abrogated the formation of HU-induced ASCC2-EdU PLA foci in both RPE1-A1 and RPE1-B1 clones (Fig. 2F and 2G). In contrast, reverting PCNA K164R back to K164K restored HU-induced ASCC2-EdU PLA foci formation in both RPE1-A1 and RPE1-B1 (Fig. 2G). These results suggest that ubiquitylated PCNA at K164 mediates recruitment of ASCC2 to stalled forks.

### ASCC2 interacts with polyubiquitylated PCNA in a manner dependent upon its ubiquitin binding activity

To gain further insight into the relationship between ASCC2 and ubiquitylated PCNA, we investigated whether ASCC2 interacts with polyubiquitylated PCNA. As we found it challenging to detect polyubiquitylated PCNA, we turned to conditions that were previously established to enable the maximal generation of PCNA polyubiquitylation^29,46^. These conditions involved UV irradiation of cells in the presence of both ATR inhibition and knockdown of USP1, which regulates PCNA ubiquitylation^47–49^, as well as formaldehyde crosslinking, which preserves potentially transient interactions of proteins with ubiquitylated PCNA. HEK293T cells transfected with the vector, HA-tagged ASCC2, or HA-tagged ASCC2 carrying LLL-AAA mutations, a ubiquitin-binding dead mutant, were exposed to these conditions, followed by coimmunoprecipitation with an anti-HA antibody. We found that HA-ASCC2 predominantly brought down polyubiquitylated PCNA whereas HA-ASCC2-LLL-AAA mutant failed to do so (Fig. 2H). These results suggest that ASCC2 relies on its ubiquitin binding activity to interact with polyubiquitylated PCNA induced by replication stress.

### ASCC3 unwinds DNA to promote RPA accumulation both in vivo and in vitro

Upon replication stress, uncoupling of leading and lagging strand DNA synthesis leads to generation of ssDNA. As ASCC3 is a DNA helicase that unwinds dsDNA with a 3’ overhang^31^, we asked whether ASCC3 plays a role in regulating the generation of ssDNA upon replication stress. To detect nascent ssDNA, we first pulse-labeled U2OS WT and ASCC3-KO cells with BrdU for 20 min prior to treatment with HU. Immunofluorescence analysis with anti-BrdU antibody under non-denaturing conditions (which only allows for detection of BrdU on ssDNA) revealed that ASCC3 loss led to a reduction in the number of cells with ≥10 BrdU foci following treatment with HU (Fig. 3A and Supplementary Fig. S4A). This reduction was unlikely due to a change in the replication fork speed, or the cell cycle profile since ASCC3 loss had little effect on either of them (Supplementary Fig. S4B-S4E). To detect parental ssDNA, we incubated U2OS WT and ASCC3-KO cells with BrdU for 20 h prior to treatment with HU. ASCC3 loss reduced the number of cells with ≥30 BrdU foci detected under non-denaturing conditions following treatment with HU (Fig. 3B). Rescue experiments revealed that while overexpression of Myc-ASCC3 restored levels of both nascent and parental ssDNA following treatment with HU in U2OS ASCC3-KO cells, overexpression of Myc-ASCC3-G1354D failed to do so (Fig. 3C and 3D). These results altogether suggest that ASCC3 uses its helicase activity to unwind DNA in response to replication stress, generating both nascent and parental ssDNA.

**Figure 3.**
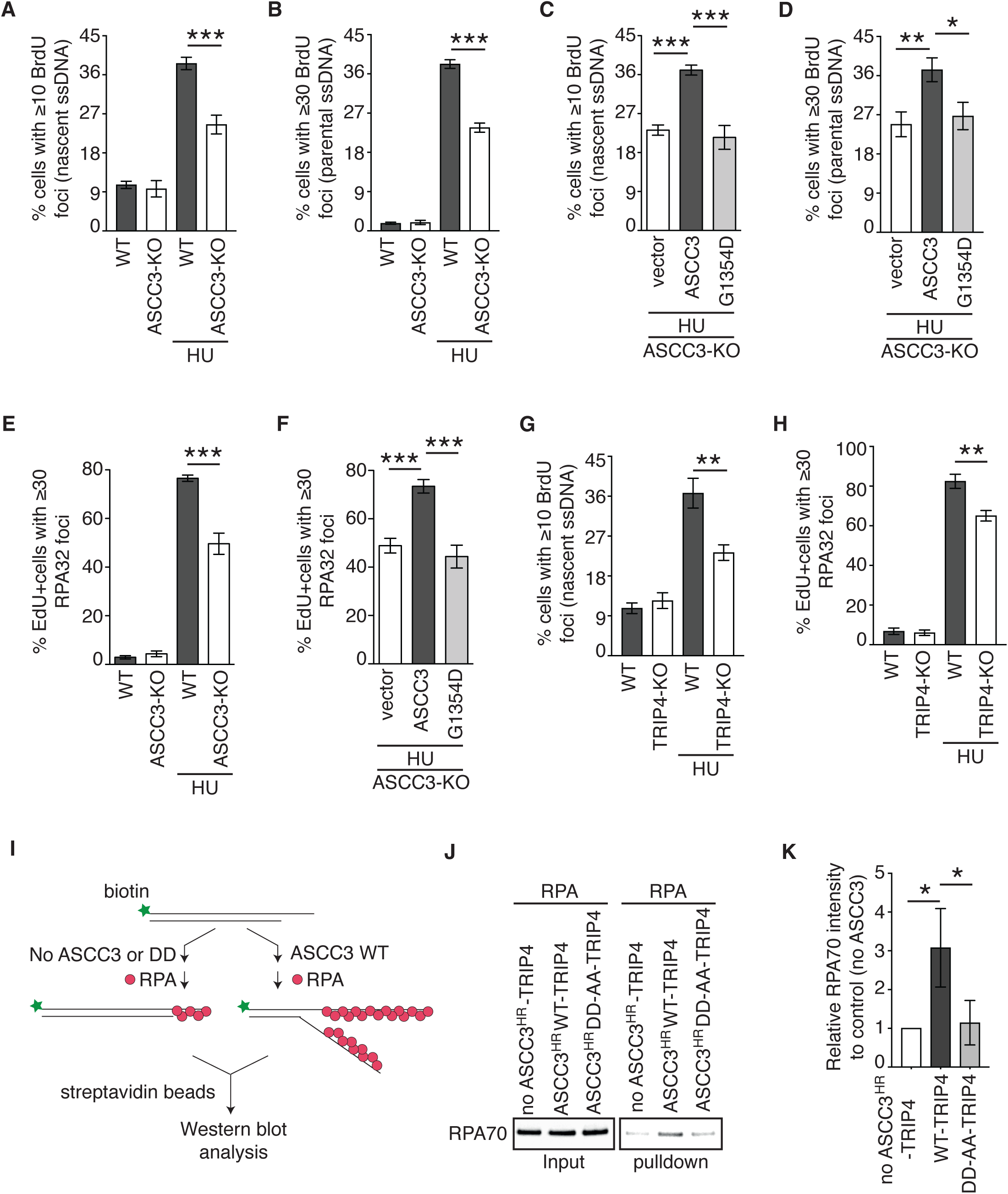
ASCC3 unwinds DNA to promote ssDNA-RPA accumulation upon replication stress. (**A**) Quantification of the percentage of cells with ≥ 10 BrdU foci in U2OS WT and ASCC3-KO cells that were pulse-labeled with BrdU for 20 min prior to their treatment with or without 4 mM HU for 4 h. A total of 802-825 cells per condition were scored in blind. SDs from three independent experiments are indicated in this and subsequent panels. \*\*\**P*<0.001. (**B**) Quantification of the percentage of cells with ≥ 30 BrdU foci in U2OS WT and ASCC3-KO cells that were labeled with BrdU for 20 h prior to their treatment with or without 4 mM HU for 4 h. A total of 800-849 cells per condition were scored in blind. \*\*\**P*<0.001. (**C**) Quantification of the percentage of cells with ≥ 10 BrdU foci in U2OS ASCC3-KO cells expressing various Myc-ASCC3 alleles as indicated. These cells were pulse-labeled with BrdU for 20 min prior to their treatment with HU. A total of 601-641 cells per condition were scored in blind. \*\*\**P*<0.001. (**D**) Quantification of the percentage of cells with ≥ 30 BrdU foci in U2OS ASCC3-KO cells expressing various Myc-ASCC3 alleles as indicated. These cells were labeled with BrdU for 20 h prior to their treatment with HU. A total of 502-521 cells per condition were scored in blind. \**P*<0.05; \*\**P*<0.01. (**E**) Quantification of EdU+ cells with ≥ 30 RPA32 foci in U2OS WT and ASCC3-KO cells that were pulse-labeled with EdU for 10 min prior to treatment with or without 4 mM HU for 6 h. A total of 800-873 cells per condition were scored in blind. \*\*\**P*<0.001. (**F**) Quantification of the percentage of EdU+ cells with ≥ 30 RPA32 foci in U2OS ASCC3-KO cells expressing various Myc-ASCC3 alleles as indicated. A total of 801-856 cells per condition were scored in blind. \*\*\**P*<0.001. (**G**) Quantification of the percentage of cells with ≥ 10 BrdU foci in U2OS WT and TRIP4-KO cells that were pulse-labeled with BrdU for 20 min prior to their treatment with or without HU. A total of 601-650 cells per condition were scored in blind. \*\**P*<0.01. (**H**) Quantification of the percentage of EdU+ cells with ≥ 30 RPA32 foci in U2OS WT and TRIP4-KO cells that were pulse-labeled with EdU for 10 min prior to their treatment with or without HU. A total of 742-977 cells per condition were scored in blind. \*\**P*<0.01. (**I**) Schematic diagram of in vitro RPA binding assays. (**J**) Western blot analyses of recombinant RPA70 recovered from streptavidin pulldown of biotinylated ssDNA from indicated reactions. Immunoblotting was done with an anti-RPA70 antibody. (**K**) Quantification of recombinant RPA70 from (J). \**P*<0.05.

RPA, a trimeric complex consisting of RPA70, RPA32, and RPA14, binds ssDNA. In agreement with the notion that ASCC3 generates ssDNA upon replication stress, we observed that ASCC3 loss also impaired the number of EdU+ cells exhibiting HU-induced RPA32 foci formation (Fig. 3E). This impairment was suppressed by Myc-ASCC3 but not Myc-ASCC3-G1354D (Fig. 3F), suggesting that ASCC3 unwinds DNA to promote efficient accumulation of RPA on ssDNA upon replication stress.

ASCC3’s helicase activity is reported to be stimulated by TRIP4^43^. To investigate whether TRIP4 regulates ASCC3 upon replication stress, we knocked out TRIP4 in U2OS cells (Supplementary Fig. S5A). While TRIP4 loss affected levels of neither ASCC2 nor ASCC3 (Supplementary Fig. S5A), it impaired the level of nascent ssDNA as evidenced by a reduction in the number of cells with ≥10 BrdU foci following treatment with HU (Fig. 3G). Loss of TRIP4 also impaired HU-induced RPA32 foci formation (Fig. 3H). To investigate whether TRIP4’s activity upon replication stress requires its interaction with ASCC3, we turned to three previously reported TRIP4 mutants that are defective in their interaction with ASCC3^43^, namely TRIP4 lacking the zinc finger domain (ΔZNF), TRIP4 carrying double CC171/184AA mutations (TRIP4-CC-AA), and TRIP4 carrying triple LLL174/180/190AAA mutations (TRIP4-LLL-AAA). Rescue experiments revealed that while overexpression of HA-TRIP4 WT restored HU-induced RPA32 foci formation in U2OS TRIP4-KO cells, overexpression of each of these three HA-tagged TRIP4 mutants failed to do so (Supplementary Fig. S5B and S5C). These results suggest that the ASCC3-TRIP4 interaction is required for generation of ssDNA upon replication stress.

To further substantiate these in vivo findings, we performed in vitro RPA binding assays in the presence or absence of recombinant ASCC3 using biotin-labeled synthetic dsDNA substrates containing a 3’ overhang (Fig. 3I). It has been reported that the helicase activity of ASCC3 is autoinhibited by its N-terminal region^42^ but stimulated by TRIP4^43^. Thus, we purified recombinant TRIP4 in complex with ASCC3 lacking the autoinhibitory N-terminal region (ASCC3^HR^-WT-TRIP4) or ASCC3^HR^ carrying helicase-dead D611A-D1453A mutations (ASCC3^HR^-DD-AA-TRIP4)^43^. The inability of ASCC3^HR^-DD-AA-TRIP4 to unwind dsDNA was also confirmed in this study (Supplementary Fig. S5D). Pulldown by streptavidin beads revealed binding of RPA to biotinylated DNA substrates with 3’ overhangs in the absence of ASCC3^HR^-TRIP4 (Fig. 3J and 3K). This binding was enhanced by ASCC3^HR^-WT-TRIP4 but not ASCC3^HR^-DD-AA-TRIP4 (Fig. 3J and 3K), supporting the notion that ASCC3 unwinds DNA to promote the RPA-ssDNA formation. Here, ASCC3^HR^-WT-TRIP4 unwound a 3’-tailed overhang duplex DNA to allow RPA binding on the ssDNA generated by its helicase activity.

### ASCC3 interacts with RPA and this interaction is further induced upon replication stress

We have shown that ASCC3 promotes RPA accumulation on ssDNA upon replication stress. RPA is found in the list of ASCC2-interacting proteins^32^ and thus, we investigated whether ASCC3 might interact with RPA. We first performed coimmunoprecipitation in HEK293T cells overexpressing GFP-ASCC3 in combination with either Myc-RPA70 or Myc-RPA32. We observed that GFP-ASCC3 was robustly brought down by Myc-RPA70 (Fig. 4A). In a reciprocal coimmunoprecipitation, Myc-ASCC3 also efficiently brought down both HA-RPA70 and HA-RPA32 (Fig. 4B and 4C). To investigate whether endogenous ASCC3 interacts with RPA, we performed coimmunoprecipitation with an anti-ASCC3 antibody. This antibody brought down both RPA70 and RPA32 (Fig. 4D). In agreement with a published report^32^, we found that an anti-ASCC2 antibody also brought down RPA70 and RPA32 (Fig. 4D). We also performed reciprocal coimmunoprecipitations with an anti-RPA32 antibody. However, we were unable to detect any significant enrichment of ASCC3 in RPA32 immunocomplexes (Fig. 4E), suggesting that it is likely that only a small fraction of endogenous RPA is engaged in its interaction with ASCC3. On the other hand, we found that both endogenous ASCC2 and ASCC3 were pulled down by exogenously expressed HA-RPA70 albeit irrespective of the presence of HU (Fig. 4F). Treatment with DNase I did not affect the ability of HA-RPA70 to bring down endogenous ASCC3 (Fig. 4G), suggesting that the interaction of ASCC3 with RPA is unlikely to be mediated by DNA. To further substantiate the interaction of ASCC3 with RPA in vivo, we performed PLA assays in both U2OS WT and ASCC3-KO cells treated with or without HU. We detected endogenous ASCC3-RPA32 PLA foci formation in unperturbed U2OS WT cells, which was further induced in HU-treated U2OS WT (Fig. 4H and 4I). Loss of ASCC3 abrogated the ASCC3-RPA32 PLA foci formation in both untreated and HU-treated cells (Fig. 4H and 4I). These results altogether suggest that ASCC3 interacts with RPA and that this interaction is further induced by replication stress.

**Figure 4.**
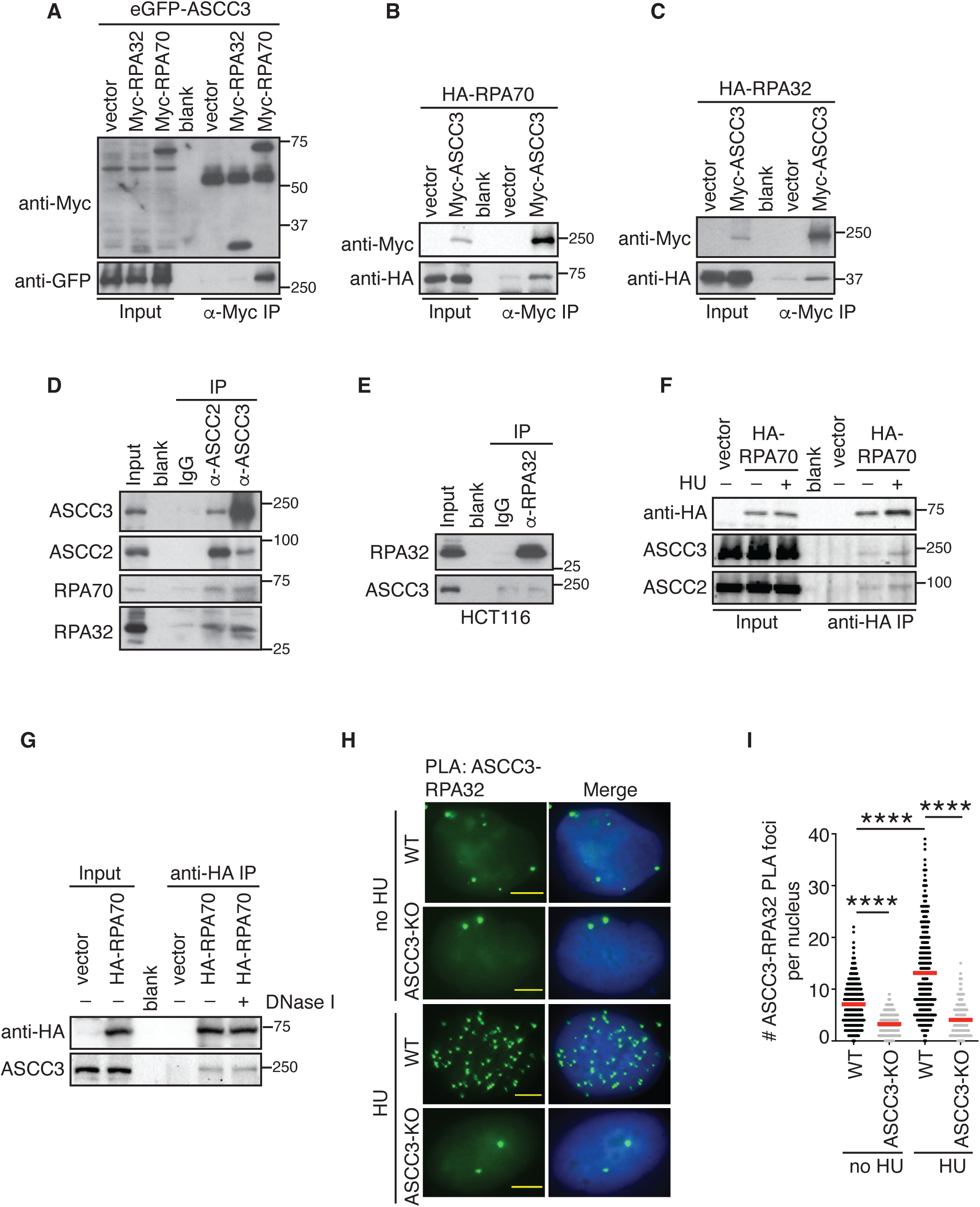
ASCC3 interacts with RPA and this interaction is further induced by replication stress. (**A**) Anti-Myc coIPs in HEK293T cells transfected with eGFP-ASCC3 together with either the vector alone, Myc-RPA32, or Myc-RPA70. Immunoblotting was done with anti-Myc and anti-GFP antibodies. (**B**) Anti-Myc coIPs in HEK293T cells transfected with HA-RPA70 together with either the vector alone or Myc-ASCC3. Immunoblotting was done with anti-Myc and anti-HA antibodies. (**C**) Anti-Myc coIPs in HEK293T cells transfected with HA-RPA32 together with the vector alone or Myc-ASCC3. Immunoblotting was done with anti-Myc and anti-HA antibodies. (**D**) Coimmunoprecipitations (coIPs) with IgG, anti-ASCC2, or anti-ASCC3 antibodies in HCT116 cells. Immunoblotting was done with anti-ASCC2, anti-ASCC3, anti-RPA70, and anti-RPA32 antibodies. (**E**) Anti-RPA32 coIPs in HCT116 cells. IgG was used as a control. Immunoblotting was performed with anti-RPA32 and anti-ASCC3 antibodies. (**F**) Anti-HA coIPs in no HU- or HU-treated HEK293T cells expressing the vector alone or HA-RPA70. Immunoblotting was done with anti-HA, anti-ASCC2, and anti-ASCC3 antibodies. (**G**) Anti-HA coIPs in HEK293T cells expressing the vector alone or HA-RPA70 in the presence or absence of DNase I. Immunoblotting was done with anti-HA and anti-ASCC3 antibodies. (**H**) Representative images of ASCC3-RPA32 PLA foci formation in no HU- or HU-treated U2OS WT and ASCC3-KO cells. (**I**) Quantification of ASCC3-RPA32 PLA foci formation from (H). The PLA experiments were performed twice independently with reproducible data. Data from one representative experiment are shown as scatter plot graphs with the mean indicated. A total of 402-418 cells per condition were analyzed. The *P*-value was determined using a non-parametric Mann-Whitney rank-sum *t*-test. \*\*\*\**P*<0.0001.

### ASCC3 promotes replication fork reversal

Native BrdU staining has been used as an indirect marker for fork reversal^18^. As we observed reduced native BrdU staining in ASCC3-KO cells (Fig. 3A), we investigated whether ASCC3 might promote fork reversal. To do so, we turned to three additional well characterized surrogate readouts for fork reversal. These readouts include 1) association of SMARCAL1 with stalled forks since it has been well documented that SMARCAL1, a chromatin remodeler, is recruited to stalled forks to catalyze fork reversal^50–52^, 2) fork progression since fork reversal is thought to restrain fork progression upon replication stress^15^, and 3) fork degradation since fork reversal has been reported to be a prerequisite for fork degradation in BRCA1/BRCA2-deficient cells^53–55^. To examine whether ASCC3 regulates recruitment of SMARCAL1 to stalled forks, we first measured HU-induced SMARCAL1 foci formation, a readout for SMARCAL1 association with stalled forks, in both U2OS WT and ASCC3-KO cells. Immunofluorescence analysis revealed that ASCC3 loss, which did not affect the level of SMARCAL1 (Supplementary Fig. S6A), led to a pronounced reduction in HU-induced SMARCAL1 foci formation (Fig. 5A). This reduction was suppressed by overexpression of Myc-ASCC3 but not Myc-ASCC3-G1354D in U2OS ASCC3-KO cells (Fig. 5B). To further substantiate this finding, we measured the SMARCAL1-EdU PLA foci formation in U2OS WT and ASCC3-KO cells that were pulse-labeled with EdU for 10 min prior to treatment with or without HU. We found that ASCC3 loss impaired HU-induced SMARCAL1-EdU PLA foci formation (Fig. 5C and 5D). This impairment was suppressed by overexpression of Myc-ASCC3 but not Myc-ASCC3-G1354D in U2OS ASCC3-KO cells (Fig. 5E). These results suggest that ASCC3 relies on its helicase activity to stimulate recruitment of SMARCAL1 to stalled forks. These results also agree with the notion that ASCC3 promotes fork reversal.

**Figure 5.**
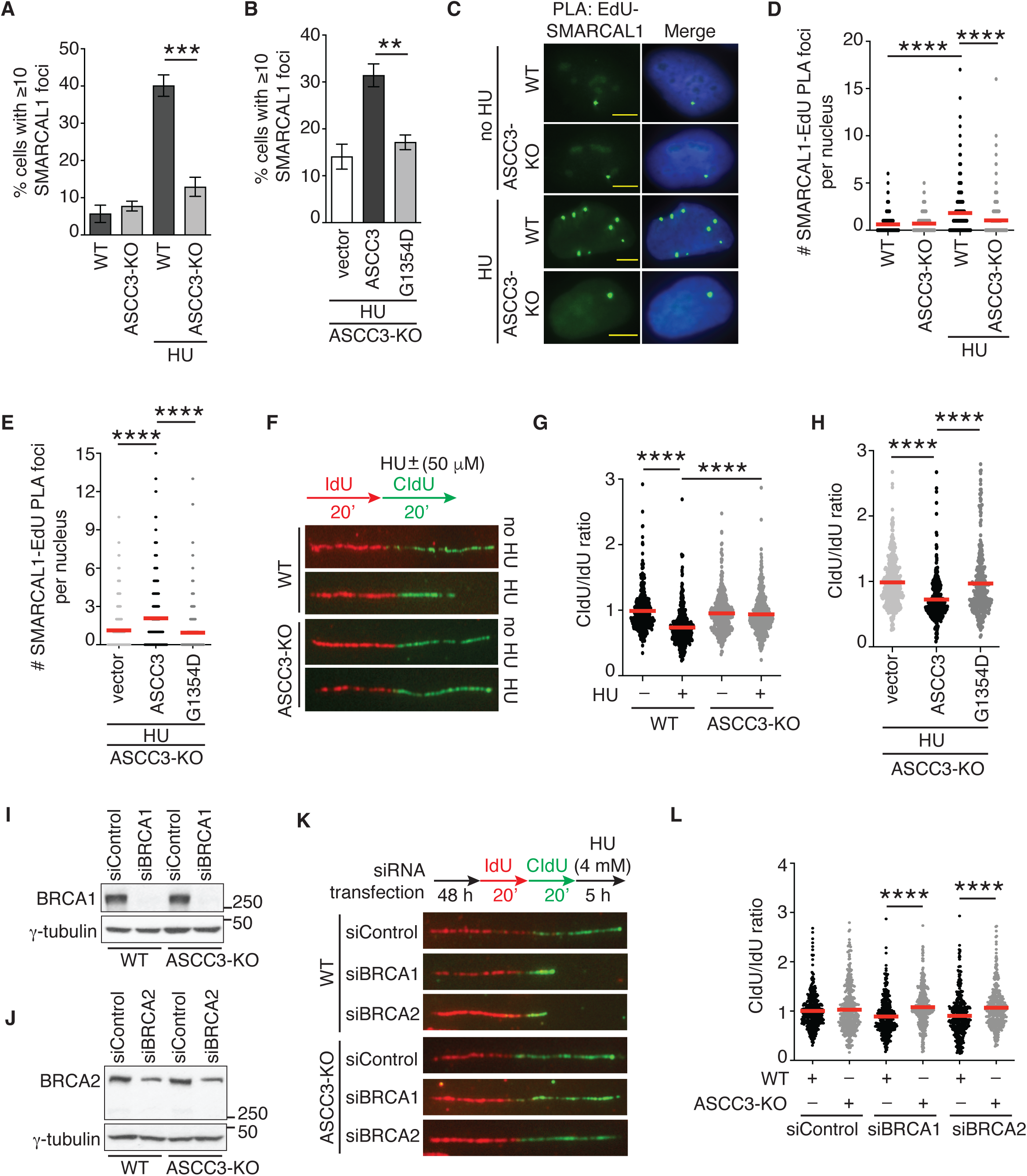
ASCC3 promotes fork reversal upon replication stress. (**A**) Quantification of the percentage of U2OS WT and ASCC3-KO cells with ≥ 10 SMARCAL1 foci. Cells were treated with or without 4 mM HU for 2 h. A total of 501-531 cells per condition were scored in blind. SDs from three independent experiments are indicated in this and 6B panels. \*\*\**P*<0.001. (**B**) Quantification of the percentage of cells with ≥ 10 SMARCAL1 foci in U2OS ASCC3-KO cells expressing the vector alone, Myc-ASCC3, or Myc-ASCC3-G1354D prior to treatment with HU. A total of 501-537 cells per condition were scored in blind. \*\**P*<0.01. (**C**) Representative images of SMARCAL1-EdU PLA foci formation in no HU- or HU-treated U2OS WT and ASCC3-KO cells. (**D**) Quantification of SMARCAL1-EdU PLA foci formation from (C). The PLA experiments were performed twice independently with reproducible data in this and 5E panels. Data from one representative experiment are shown as scatter plot graphs with the mean indicated in this and subsequent panels. A total of 768-793 cells per condition were analyzed. The *P*-value was determined using a non-parametric Mann-Whitney rank-sum *t*-test in this and subsequent panels. \*\*\*\**P*<0.0001. (**E**) Quantification of SMARCAL1-EdU PLA foci formation in HU-treated U2OS ASCC3-KO cells expressing the vector alone, Myc-ASCC3, or Myc-ASCC3-G1354D. A total of 612-635 cells per condition were analyzed. \*\*\*\**P*<0.0001. (**F**) Representative images of DNA fibers from U2OS WT or ASCC3-KO cells treated with or without HU. (**G**) Quantification of the CldU/IdU ratio from (F). DNA fiber experiments were performed three times independently with reproducible data. A total of 388-418 fibers per condition were analyzed. \*\*\*\**P*<0.0001. (**H**) Quantification of the CldU/IdU ratio for HU-treated U2OS ASCC3-KO cells expressing the vector alone, Myc-ASCC3, or Myc-ASCC3-G1354D. DNA fiber experiments were performed twice independently with reproducible data. A total of 333-357 fiber per condition were analyzed. \*\*\*\**P*<0.0001. (**I**) Western blot analyses of U2OS WT and ASCC3-KO cells transfected with indicated siRNAs. Immunoblotting was performed with anti-BRCA1 and anti-γ-tubulin antibodies. (**J**) Western blot analysis of U2OS WT and ASCC3-KO cells transfected with indicated siRNA. Immunoblotting was performed with anti-BRCA2 and anti-γ-tubulin antibodies. (**K**) Representative images of DNA fibers from U2OS WT or ASCC3-KO cells transfected with siControl, siBRCA1, or siBRCA2. (**L**) Quantification of the CldU/IdU ratio from (K). DNA fiber experiments were performed twice independently with reproducible data. A total of 400-407 fibers per condition were analyzed. \*\*\*\**P*<0.0001.

To investigate whether ASCC3 restrains fork progression upon replication stress, we pulse-labeled U2OS WT and ASCC3-KO cells first with IdU for 20 min and then with CldU for 20 min in the presence of a low dose of HU (50 μM). DNA fiber analysis revealed that ASCC3 loss restored the ratio of CldU/IdU in the presence of 50 μM HU in U2OS cells (Fig. 5F and 5G). This restoration in the ratio of CldU/IdU was also observed in HCT116 cells in which ASCC3 had been either knocked out (Supplementary Fig. S6B and S6C) or knocked down for ASCC3 (Supplementary Fig. S6D and S6E). To further substantiate this finding, we performed rescue experiments in U2OS ASCC3-KO cells. While overexpression of Myc-ASCC3 reduced the ratio of CldU/IdU in the presence of 50 μM HU in ASCC3-KO cells, overexpression of Myc-ASCC3 carrying the helicase-dead G1354D mutation had little effect on the ratio of CldU/IdU in the presence of 50 μM HU in ASCC3-KO cells (Fig. 5H). These results suggest that ASCC3 requires its helicase activity to restrain fork progression in response to mild replication stress. As fork reversal can slow down fork progression^15^, these results lend support to the notion that ASCC3 promotes fork reversal.

To investigate whether ASCC3 promotes fork degradation in BRCA1/BRCA2-deficient cells, we knocked down BRCA1 or BRCA2 in both U2OS WT and ASCC3-KO cells (Fig. 5I and 5J). These cells were first labeled with IdU and then CldU prior to treatment with 4 mM HU for 5 h. DNA fiber analysis revealed that while depletion of either BRCA1 or BRCA2 led to a reduction in the ratio of CldU/IdU, this reduction was suppressed by loss of ASCC3 (Fig. 5K and 5L). These results fall in line with the notion that loss of fork reversal in ASCC3-KO cells restores fork stability in the absence of BRCA1/BRCA2, lending further support to the idea that ASCC3 promotes fork reversal.

### ASCC3 inhibits PRIMPOL- and RAD51-mediated damage tolerance pathways to restrain fork progression upon replication stress

Upon replication stress, DNA damage tolerance pathways can be engaged to support fork progression. To investigate whether ASCC3 inhibits DNA damage tolerance pathways to restrain fork progression upon replication stress, we first knocked down PRIMPOL (Supplementary Fig. S7A), which is responsible for fork repriming, in U2OS WT and ASCC3-KO cells. DNA fiber analysis revealed that depletion of PRIMPOL abrogated restoration of the ratio of CldU/IdU in ASCC3-KO cells in the presence of 50 μM HU (Fig. 6A and Supplementary Fig. S7B). PRIMPOL-dependent fork repriming is associated with accumulation of ssDNA gaps^11,56,57^. In agreement with previous reports, the restored ratio of CldU/IdU in ASCC3-KO cells in the presence of 50 μM HU was sensitive to S1 nuclease (Supplementary Fig. S7C and S7D). These results suggest that ASCC3 restrains fork progression by inhibiting PRIMPOL-dependent fork repriming upon replication stress.

**Figure 6.**
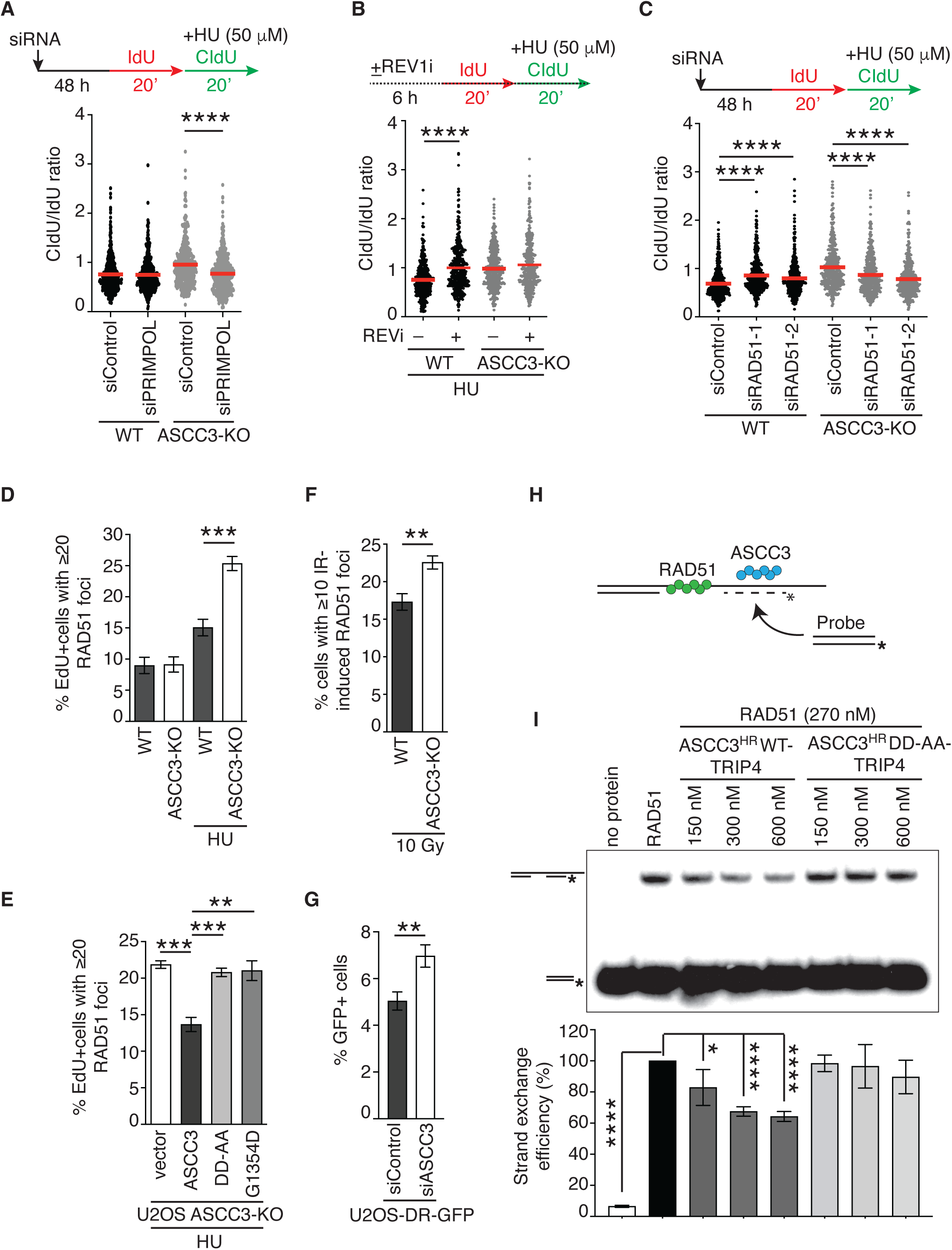
ASCC3 antagonizes RAD51-mediated recombination. (**A**) Quantification of the CldU/IdU ratio in HU-treated U2OS WT and ASCC3-KO cells expressing control siRNA or siRNA against PRIMPOL (siPRIMPOL). DNA fiber experiments were performed twice independently with reproducible data in this, 6B, and 6C panels. Data from one representative experiment are shown as scatter plot graphs with the mean indicated in this, 6B, and 6C panels. A total of 407-421 fibers per condition were analyzed. The *P*-value was determined using a non-parametric Mann-Whitney rank-sum *t*-test in this and subsequent panels. \*\*\*\**P*<0.0001. (**B**) Quantification of the CldU/IdU ratio in U2OS WT and ASCC3-KO cells that were treated with HU in the presence or absence of REV1 inhibitor JH-RE-06 (REV1i). A total of 413-414 fibers per condition were analyzed. \*\*\*\**P*<0.0001. (**C**) Quantification of the CldU/IdU ratio in HU-treated U2OS WT and ASCC3-KO cells expressing control siRNA or two independent siRNA against RAD51 as indicated. A total of 407-415 fibers per condition were analyzed. \*\*\*\**P*<0.0001. (**D**) Quantification of EdU+ cells with ≥ 20 RAD51 foci in U2OS WT and ASCC3-KO cells that were pulse-labeled with EdU for 10 min prior to treatment with or without 4 mM HU for 6 h. A total of 801-853 cells per condition were scored in blind. SDs from three independent experiments are indicated in this, and 6E-6G panels. \*\*\**P*<0.001. (**E**) Quantification of EdU+ cells with ≥ 20 RAD51 foci in U2OS ASCC3-KO cells expressing various Myc-ASCC3 alleles as indicated. A total of 802-827 cells per condition were scored in blind. \*\**P*<0.01; \*\*\**P*<0.001. (**F**) Quantification of cells with ≥ 10 IR-induced RAD51 foci in U2OS WT and ASCC3-KO cells that were treated with 10 Gy IR prior to fixation. A total of 503-554 cells per condition were scored in blind. \*\**P*<0.01. (**G**) Quantification of the percentage of cells with restoration of GFP expression following HR-mediated repair of I-SceI-induced DSBs. \*\**P*<0.01. (**H**) Schematic diagram of strand exchange assays. (**I**) In vitro strand exchange assays. Top panel: Recombinant RAD51 (270 nM) was premixed with 1µM of 3’-ssDNA tail for 5 min, followed by further incubation with 350 nM of labeled duplex DNA and increasing concentration (150 nM, 300 nM, 600 nM) of ASCC3^HR^ WT-TRIP4 or ASCC3^HR^-DD-AA-TRIP4 for 15 min. * indicates the ^32^P-labeled DNA end. Bottom panel: Quantification of strand exchange efficiency (%) relative to RAD51 alone reaction from top panel. SDs from four independent experiments are shown. \**P*<0.05; \*\*\*\**P*<0.0001.

To investigate whether translesion synthesis (TLS) and template switching (TS) mediate unrestrained fork progression in ASCC3-KO cells upon replication stress, we either inhibited REV1, a DNA polymerase involved in TLS^56^ or knocked down RAD51, a key factor in TS, in both U2OS WT and ASCC3-KO cells. We found that treatment with the REV1 inhibitor JH-RE-06 had little effect on the ratio of CldU/IdU in the presence of 50 μM HU in ASCC3-KO cells (Fig. 6B and Supplementary Fig. S7E), suggesting that in the absence of ASCC3, unrestrained fork progression does not involve TLS. In contrast, depletion of RAD51 with two independent siRNAs impaired restoration of the ratio of CldU/IdU in the presence of 50 μM HU in ASCC3-KO cells (Fig. 6C; Supplementary Fig. S7F and S7G), suggesting that in the absence of ASCC3, unrestrained fork progression is mediated by the RAD51-dependent damage tolerance pathway.

### ASCC3 functions to antagonize RAD51-dependent recombination both in vivo and in vitro

To further investigate how ASCC3 inhibits RAD51 upon replication stress, we measured HU-induced RAD51 foci formation, a readout for RAD51 association with stalled forks, in both U2OS WT and ASCC3-KO cells. We found that ASCC3 loss stimulated HU-induced RAD51 foci formation (Fig. 6D). This stimulation was suppressed by overexpression of Myc-ASCC3 but not two helicase-dead Myc-ASCC3 mutants, Myc-ASCC3-G1354D and Myc-ASCC3-D611A-D1453A (Myc-ASCC3-DD-AA) in U2OS ASCC3-KO cells (Fig. 6E), suggesting that ASCC3 requires its helicase activity to inhibit excessive RAD51 recruitment to stalled forks.

To investigate whether ASCC3 inhibits RAD51-dependent homologous recombination (HR), we first evaluated the formation of ionizing radiation (IR)-induced RAD51 foci, readouts for HR activity, in both U2OS WT and ASCC3-KO cells. ASCC3 loss increased IR-induced RAD51 foci formation (Fig. 6F). To substantiate this finding, we turned to a previously described U2OS DR-GFP reporter cell line^58,59^, in which restoration of GFP expression is dependent upon RAD51-mediated HR of an I-SceI-induced DSB in the GFP gene. Using this cell line, depletion of ASCC3 enhanced HR-mediated restoration of GFP expression (Fig. 6G). As RAD51-mediated strand exchange is a key step in HR, we further investigated whether ASCC3 inhibits this step using recombinant proteins RAD51 and ASCC3 as well as synthetic DNA substrates (Fig. 6H). To do so, we again turned to ASCC3^HR^-WT-TRIP4 and ASCC3^HR^-DD-AA-TRIP4. In vitro strand exchange assays revealed that ASCC3^HR^ WT-TRIP4 but not ASCC3^HR^-DD-AA-TRIP4 was able to inhibit RAD51-catalyzed strand exchange (Fig. 6I), suggesting that ASCC3 possesses an anti-recombinase activity.

### ASCC3 promotes efficient ATR activation and genomic stability upon replication stress

RPA bound to ssDNA is a trigger for activation of ATR, which directly phosphorylates RPA32 on S33 and CHK1 on S345^5^. As we have shown that ASCC3 stimulates the formation of ssDNA-RPA upon replication stress, we asked whether ASCC3 regulates ATR activation upon replication stress. To address this question, we measured levels of RPA32-pS33 and CHK1-pS345 in both U2OS and HCT116 WT and ASCC3-KO cells following treatment with HU. Western analysis revealed that ASCC3 loss severely impaired the level of HU-induced RPA32-pS33 in both U2OS and HCT116 cells (Fig. 7A). ASCC3 loss also led to a decrease in the level of HU-induced CHK1-pS345 in both U2OS and HCT116 cells (Fig. 7A). Aside from HU, we also investigated whether ASCC3 regulates ATR activation in response to other replication stress-inducing agents such as UV, camptothecin (CPT), and aphidicolin. We observed that ASCC3 loss led to a pronounced reduction in the level of UV-induced RPA32-pS33 (Fig. 7B). ASCC3 loss also impaired the level of CPT-induced RPA32-pS33 (Fig. 7B). We noticed that aphidicolin was a poor inducer of RPA32-pS33 (Fig. 7B), in agreement with a previous report^60^. On the other hand, UV, CPT, and aphidicolin were all able to induce CHK1-pS345 in U2OS WT cells, which was abrogated in U2OS ASCC3-KO cells (Fig. 7B). These results altogether suggest that ASCC3 promotes efficient ATR activation upon replication stress.

**Figure 7.**
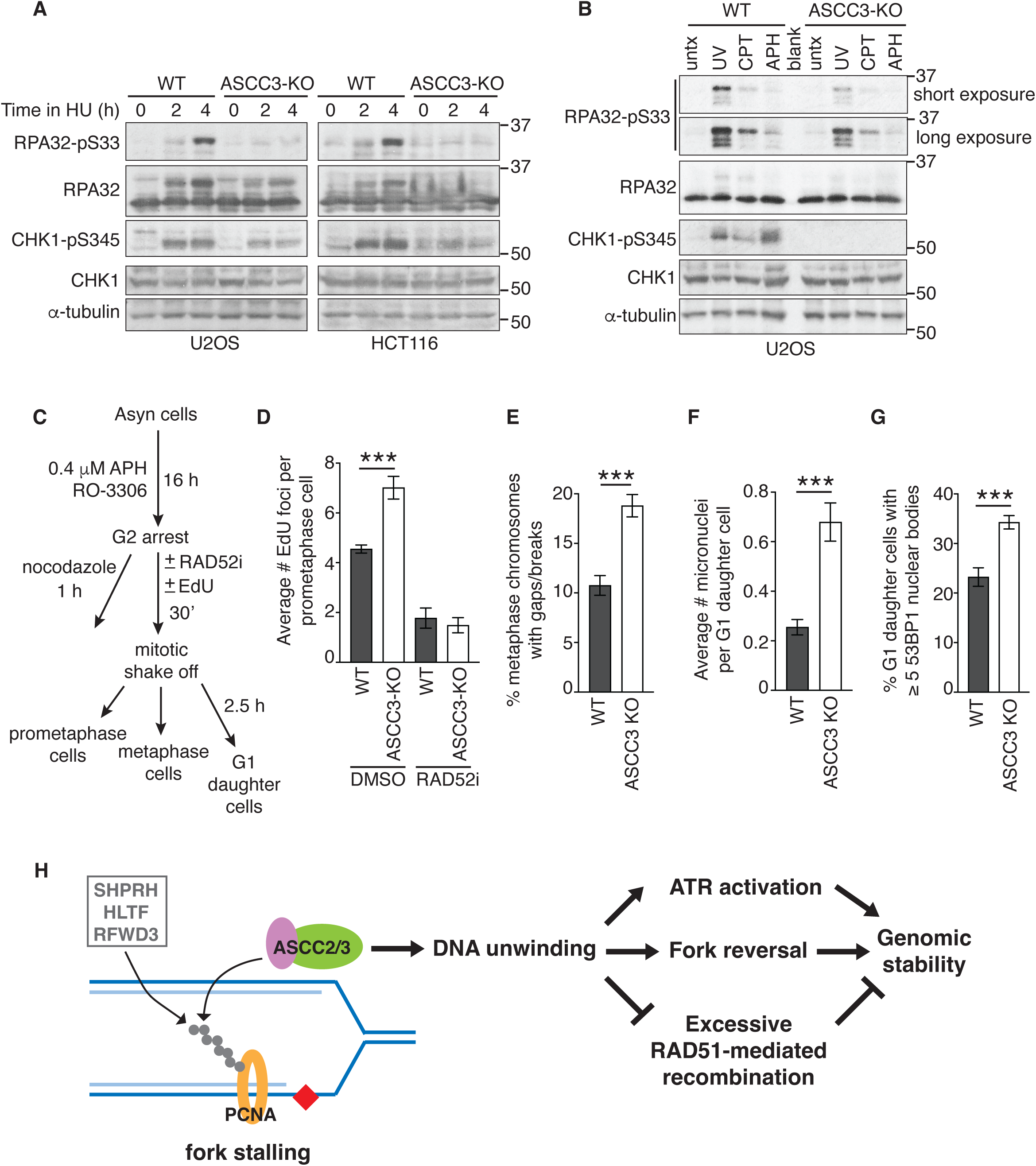
ASCC3 promotes efficient ATR activation and genomic stability. (**A**) Western blot analyses of U2OS and HCT116 WT and ASCC3-KO cells treated with no HU or HU (4 mM) for various times as indicated. Immunoblotting was done with antibodies against RPA32-pS33, RPA32, CHK1-pS345, CHK1, and α-tubulin. (**B**) Western blot analyses. U2OS WT and ASCC3-KO cells were either untreated or treated with UV (30 J/m^2^), camptothecin (CPT) (1 μM CPT for 1 h) or aphidicolin (APH) (0.4 μM for 24 h). Immunoblotting was done with antibodies against RPA32-pS33, RPA32, CHK1-pS345, CHK1, and α-tubulin. (**C**) Schematic diagram of experimental setup. APH: aphidicolin; RO-3306: CDK1 inhibitor; RAD52i: RAD52 inhibitor AICAR. (**D**) Quantification of the average number of EdU foci in both U2OS WT and ASCC3-KO prometaphase cells treated with or without RAD52i (20 μM). A total of 188-207 prometaphase cells per condition were scored in blind. SDs from three independent experiments are shown in this and subsequent panels. \*\*\**P*<0.001. (**E**) Quantification of the percentage of metaphase chromosomes with gaps and breaks in both U2OS WT and ASCC3-KO following exposure to aphidicolin in S phase. A total of 20 metaphase cells (with a total of 905-975 metaphase chromosomes) per condition were scored in blind. \*\*\**P*<0.001. (**F**) Quantification of the average number of micronuclei in both U2OS WT and ASCC3-KO G1 daughter cells following exposure to aphidicolin in previous S phase. A total of 508-553 cells per condition were scored in blind. \*\*\**P*<0.001. (**G**) Quantification of the percentage of both U2OS WT and ASCC3-KO G1 daughter cells with ≥ 5 53BP1 nuclear bodies following exposure to aphidicolin in previous S phase. A total of 510-517 cells per condition were scored in blind. \*\*\**P*<0.001. (**H**) Model for control of replication stress responses by ASCC3. See the text for details.

Impaired ATR activation can lead to under-replicated DNA in S phase, which can be repaired through a form of RAD52-dependent break-induced replication^61,62^, also known as mitotic DNA synthesis (MiDAS), throughout G2 and into early mitosis. To investigate whether impaired ATR activation observed in the absence of ASCC3 is associated with elevated MiDAS activity, we treated U2OS WT and ASCC3-KO cells with aphidicolin in the presence of CDK1 inhibitor RO3306 for 16 h (Fig. 7C). RO3306 arrested these cells in G2. Following their release from G2, these cells were pulse-labeled with EdU in early mitosis. ASCC3 loss was found to increase incorporation of EdU in prometaphase cells, and this increase was sensitive to treatment with the RAD52 inhibitor AICAR (Fig. 7D and Supplementary Fig. S8A). These results agree with the notion that ASCC3 promotes ATR activation to prevent DNA under-replication in S phase. As under-replicated DNA can drive genomic instability, we further evaluated frequencies of chromosome breaks and mis-segregation in ASCC3-KO cells. We found that ASCC3 loss induced the formation of chromatid breaks/gaps, micronuclei and 53BP1 nuclear bodies in G1 daughter cells following exposure to aphidicolin in S phase (Fig. 7E-7G; Supplementary Fig. S8B-S8D). These results altogether suggest that ASCC3 promotes genomic stability upon replication stress.

## DISCUSSION

The Ski2-like helicase ASCC3 participates in a diverse set of molecular processes ranging from transcriptional regulation, alkylation damage repair, RNA splicing to ribosome quality control^30–35^. However, little is known about whether it plays a role in the cellular response to replication stress. The work presented in this report has uncovered that ASCC3 is recruited to stalled forks by its interacting partner ASCC2 and controls multiple replication stress responses, including fork reversal, ATR activation, and RAD51-dependent recombination (Fig. 7H).

Several lines of evidence presented here suggest that ASCC2 recruits ASCC3 to stalled forks by binding PCNA polyubiquitylated at K164 catalyzed by SHPRH, HLTF, and RFWD3 ligases (Fig. 7H). Firstly, ASCC2 relies on both its ubiquitin binding activity and ubiquitylation of PCNA at K164 to be recruited to stalled forks. Secondly, recruitment of ASCC2 to stalled forks is dependent upon SHPRH, HLTF, and RFWD3, E3 ligases that polyubiquitylate PCNA at K164^25–27^. Thirdly, ASCC2 predominantly pulls down polyubiquitylated PCNA upon replication stress in a manner dependent upon its ubiquitin binding activity. Although these findings do not rule out the possibility that monoubiquitylation of PCNA recruits ASCC2 to stalled forks, our finding that ASCC3 inhibits RAD51-but not REV1-mediated damage tolerance pathway in the presence of mild replication stress (50 μM HU) (Fig. 6B and 6C) argues against this possibility since polyubiquitylation of PCNA at K164 channels the damage tolerance pathway to RAD51-mediated template switching^25–27^.

Our work suggests that upon replication stress, unwinding of DNA by ASCC3 is a prerequisite to fork reversal (Supplementary Fig. S9). In support of this model, aside from impaired generation of ssDNA upon replication stress, loss of ASCC3 leads to 1) defective SMARCAL1 recruitment to stalled forks, 2) reduced native BrdU staining, 3) unrestrained fork progression upon mild replication stress, and 4) restored fork stability in BRCA1/2-deficient cells in response to fork stalling, all of which are readouts for fork reversal. Furthermore, ASCC3 loss engages PRIMPOL-mediated fork repriming, which is inhibited by fork reversal^13,14^, to support unrestrained fork progression upon mild replication stress, agreeing with the notion that ASCC3 promotes fork reversal to inhibit PRIMPOL-mediated fork repriming (Supplementary Fig. S9). We have shown that a helicase-dead ASCC3-G1354D mutant fails to promote SMARCAL1 recruitment to stalled forks (Fig. 5E), suggesting that ASCC3 unwinds DNA upstream of SMARCAL1 recruitment upon replication stress. Fork reversal involves annealing of nascent DNA strands, converting a three-way junction to a four-way junction^5,6,18^. Recently, it has been suggested that this process might involve unwinding of nascent strands from parental DNA and then their direct annealing to each other^17^. Although which helicases are involved in this process is unknown, it has been speculated that FBH1 might perform the unwinding of nascent strands in this model^17^. Our finding suggests that ASCC3 is also a helicase that could fulfill this function (Supplementary Fig. S9), lending support to the idea that active unwinding of nascent strands is required for fork reversal.

We have shown that ASCC3 interacts with RPA and that this interaction is unlikely to be mediated by DNA. While we cannot rule out the possibility that ASCC3 interacts with RPA indirectly, our work suggests that it is likely that in addition to generating ssDNA for RPA, ASCC3 also interacts with RPA to promote RPA accumulation on ssDNA in response to replication stress. As it has been well documented that RPA bound ssDNA is a trigger for ATR activation^5,8^, our work suggests that ASCC3 promotes efficient ATR activation by stimulating RPA accumulation on ssDNA upon replication stress (Supplementary Fig. S9).

Deregulated RAD51-mediated recombination contributes to genomic instability^63–65^. Helicases such as FBH1, RTEL1, and BLM have been previously reported to have anti-recombinase activities^20,66–68^. Our work presented here suggests that ASCC3 also functions as an anti-recombinase since 1) loss of ASCC3 increases the formation of IR-induced RAD51 foci, readouts of HR; 2) loss of ASCC3 enhances HR-dependent restoration of GFP expression in a reporter assay; and 3) ASCC3 inhibits RAD51-mediated strand exchange in vitro. Aside from HR, RAD51 also promotes fork reversal independently of its strand exchange activity^69^, key to HR, as well as protects nascent DNA in the reversed arm following fork reversal^70,71^. We have shown that RAD51 mediates unrestrained fork progression in ASCC3-KO cells. As there is loss of fork reversal in these cells, this activity of RAD51 is separate from RAD51’s role in fork reversal, or its protective function of reversed strands. Instead, our data suggest that in the absence of ASCC3, RAD51-dependent HR activity, likely template switching, supports unrestrained fork progression in ASCC3-KO cells upon replication stress. Our work suggests that ASCC3 prevents excessive DNA recombination upon replication stress (Supplementary Fig. S9).

Unregulated fork reversal, compromised ATR activation, and excess DNA recombination can all lead to genomic instability^63–65,72,73^. Our finding that loss of ASCC3 induces both chromosome breaks/gaps and mis-segregation underscores a previously uncharacterized but critical role of the Ski2-like helicase ASCC3 in controlling multiple replication stress responses to maintain genomic stability (Fig. 7H).

## MATERIALS AND METHODS

### Plasmids, siRNA, antibodies and drugs

Retroviral expression constructs pLPC-N-Myc-ASCC2, pLPC-N-Myc-ASCC2 carrying triple L^478^L^479^P^480^-AAA mutations, and pLPC-N-Myc-ASCC3 were respectively derived from pCMV6-MYKDDK-ASCC2^33^, pCMV6-MYKDDK-ASCC2-mut1^33^, and pcDNA3.1-ASCC3-3xFLAG^33^, all of which were kind gifts from Ramanujan Hegde. The pLPC-N-Myc-ASCC3 plasmid was used as a template to generate, via site-directed mutagenesis, pLPC-N-Myc-ASCC3 carrying either a G1345D mutation or double D611A-D1453A (DD-AA) mutations. TRIP4 expression constructs (pHAGE-HA-TRIP4 WT, pHAGE-HA-TRIP4-ΔZNF, pHAGE-HA-TRIP4-C171A-C184A, and pHAGE-HA-TRIP4-L174A-L180A-L190A) (generous gifts from Nima Mosammaparast) have been described^43^. RPA expression constructs (pLPC-NMyc-RPA70, pLPC-NMyc-RPA32, pLPC-NFH2-RPA70 and pLPC-NFH2-RPA32) have been described^74^. The cDNA of SHPRH, which was from mammalian gene collection, was used to generate the retroviral expression construct pLPC-N-Myc-SHPRH, which was subsequently used as a template to derive, via site-directed mutagenesis, pLPC-N-Myc-SHPRH carrying a C1432A mutation. All constructs were verified by Sanger sequencing.

siRNAs used were from Dharmacon: non-targeting siRNA (siControl; D-001206-14-05); siASCC2 (GGUAUUUUGUGUUAUACAA); siASCC3 (GCAAGAUAAUUAUAAUGAA); siBRCA1 (D-003461-05); siBRCA2 (D-003462-04); siCSB (GAAGAGUUGUCAGUGAUUA); siHLTF (GGAAUAUAAUGUUAACGAU); siPRIMPOL (GAGGAAAGCUGGACAUCGA); siRAD51-1 (AAGGGAAUUAGUGAAGCCAAA); siRAD51-2 (GAAGCUAUGUUCGCCAUUA); siRFWD3 (GGACCUACUUGCAAACUAU); siSHPRH (GCACAAAUCAGUCGUGUUA); siSMARCAL1 (GAAUCUCACUUCCUCAAAA); siTRIP4 (CUUCAAAAAAGAUGAAAUU); siUSP1 (GAAAUACACAGCCAAGUAA). Drugs used were aphidicolin (Sigma); camptothecin (Sigma); hydroxyurea (BioShop Canada); RAD52 inhibitor (AICAR, Cayman Chemical); REV1 inhibitor JH-RE-06 (SML2993, Sigma); RO-3306 (Selleck Chemicals).

Antibodies used include: ASCC2 (1:1000; A304-020A, Bethyl Laboratories); ASCC2 (1:5000; 1152-1-AP, Proteintech); ASCC3 (1:400; 17627-1-AP, Proteintech); ASCC3 (1:1000; PA5-56794, Invitrogen); Biotin (1:100000; A150-109A, Bethyl Laboratory); Biotin (1:100000; 200-002-211, Jackson ImmunoResearch); BrdU (1:100; 347580, BD Biosciences); BrdU (1:500; MAB3222, Millipore); BrdU (BU1/75 [ICR1]) (1:800; NB500-169, Novus Biologicals); 53BP1 (1:2000; 612522, BD Biosciences); BRCA1 (1:5000; 07-434, Millipore); BRCA2 (1:2000; 29450-1-AP, Proteintech); CHK1 (1:250; sc-7898, Santa cruz); CHK1-pS345 (1:1000; 2348S, Cell Signaling); CSB (1:200; ab66598, Abcam); FASN (1:20000; 10624-2-AP, Proteintech); GFP (1:1000; 50430-2-AP, Proteintech); HA (1:500; #2367, Cell Signaling); HLTF (1:1000; A300-229A, Bethyl Laboratory); Myc (1:1000; 9E10, Calbiochem); PCNA (1:2000; sc-56, Santa Cruz); PCNA (1:600 for PLA or 1:4000 for western blot; 10205-2-AP, Proteintech); Ubiquityl-PCNA (K164) (1:1000; 13439T, Cell Signaling); PRIMPOL (1:1000; 29824-1-AP, Proteintech); RAD51 (1:2000, ab63801, Abcam); RFWD3 (1:1000; 19893-1-AP, Proteintech); RPA32 (1:10000; NB100-332, Novus Biologicals); RPA32 (1:200; ab2175, Abcam); RPA32-pS33 (1:50000; A300-246A, Bethyl Laboratories); RPA70 (1:2000; 2267S, Cell Signaling); SHPRH (21995-1-AP, Proteintech); SMARCAL1 (1:100, sc-166209, Santa cruz); SMARCAL1 (1:2000; GTX109468, GenTex); TRIP4 (1:1000; 12324-1-AP, Proteintech); α-tubulin (1:20,000; T9026, Sigma); γ-tubulin (1:20,000; GTU88, Sigma).

### Cell culture, transfection, retroviral infection

All cells were grown in DMEM medium with 10% fetal bovine serum supplemented with non-essential amino acids, L-glutamine, 100 U/ml penicillin and 0.1 mg/ml streptomycin. Cell lines used: RPE-1 parental^45^, RPE-1 A1 (*PCNA*^K164R^)^45^; RPE-1 B1 (*PCNA*^K164R^)^45^, RPE-1 A1 revertant (*PCNA*^K164K^)^45^, RPE-1 B1 revertant (*PCNA*^K164K^)^45^, U2OS (ATCC), U2OS-DR-GFP^58^, HCT116 (Life Technology), HEK293T (ATCC), and Phoenix^58^ (a kind gift from Titia de Lange). Cell cultures were routinely fixed, stained with DAPI, and examined for mycoplasma contamination. Retroviral gene delivery was carried out as described^58^ to generate stable cell lines. DNA and siRNA transfections were carried out with Lipofectamine 2000 (Invitrogen) and RNAiMAX (Invitrogen) respectively according to the manufacturer’s instructions.

### CRISPR/Cas9 genome editing

Cells were transiently transfected with gRNA that was cloned into and expressed from the pX459v2 (#62988, Addgene) containing Cas9 followed by 2A-Puromycin cassette. The gRNA sequences are: ASCC2, 5’-GCCAAGTTACTACAGTGACC-3’; ASCC3, 5’-GACATTTGAAAAGGAACGCA-3’; TRIP4, 5’-AAAGGTGGACATCTCTACCA-3’; SHPRH, 5’-TTGTGACAAGGGTATTCTGG-3’. Following selection with puromycin for 2 days, cells were subcloned to allow for the formation of single colonies. Individual clones were confirmed by western blot analyses with anti-ASCC2, anti-ASCC3, anti-TRIP4, or anti-SHPRH antibodies for their respective loss.

### PLA assays

PLA assays were performed as described^74,75^ using Duolink® PLA kit (Sigma) according to the manufacturer’s instructions. Briefly, coverslips were blocked in Duolink® blocking solution for 30 min at 37°C and then incubated with primary antibody diluted in Duolink® antibody diluent overnight at 4°C. Subsequently, coverslips were washed twice in wash buffer A [0.15 M NaCl, 10 mM Tris-HCl (pH7.4), 0.05% Tween-20] for 5 min and then incubated with anti-rabbit PLUS and anti-mouse MINUS PLA probes diluted in Duolink® antibody diluent for 1 h at 37°C. Following washes twice in Wash Buffer A for 5 min, coverslips were subject to ligation reactions for 30 min at 37°C. Coverslips were then washed twice in Wash Buffer A for 5 min. Following amplification that was performed using Duolink® *In Situ* Detection Reagents Green for 100 min at 37°C, coverslips were washed twice in Wash Buffer B (0.1 M NaCl, 0.2 M Tris) for 10 min and once in 0.1x Wash Buffer B for 1 min. Lastly, coverslips were stained with DAPI (100 ng/ml in PBS). Cell images were captured on a Zeiss Axio Imager M2 microscope with a Zeiss Axiocam 705 mono camera and processed using Zeiss ZEN lite software. PLA signals were quantified in ImageJ (NIH) with these settings: the Wand Tool (Legacy mode) with a tolerance score of 20 and the “Smooth if thresholded” option to define individual nuclei, and the “Find Maxima” function with a prominence score set above 15.0. These settings were applied to all images. PLA data were plotted using PRISM (version 10.1.1). For scatter plot appearance, the “Standard” option was chosen as recommended by PRISM.

### DNA fiber analysis

DNA fiber analysis was done essentially as described^74,76^. For fork protection, cells were incubated first with 25 μM IdU (I7125, Sigma) for 20 min and then 250 μM CldU (C6891, Sigma) for 20 min prior to treatment with 4 mM HU for 5 h. For fork progression, cells were incubated first with 25 μM IdU for 20 min and then 250 μM CldU for 20 min in the presence of 50 μM HU. Following labeling, cells were collected by trypsinization and counted. Cells were then spotted onto one end of a glass slide, lysed in freshly made lysis buffer (50 mM EDTA pH 8.0, 200 mM Tris-HCl pH 7.5, 0.5% SDS) for 5 min, and stretched onto the slide. Slides were then fixed in freshly made methanol:acetic acid (3:1) for 20 min at −20°C and then allowed to air dry. Following incubation in freshly prepared 2.5 M HCl for 80 min, slides were washed three times in PBS and blocked with 5% BSA in PBS for 20 min at room temperature. Slides were then incubated with both rat anti-BrdU (1:800, NB500-169, Novus Biologicals) and mouse anti-BrdU (1:100, 347580, BD Sciences) antibodies prepared in 5% BSA in PBS for overnight at 4°C. Subsequently, slides were washed three times in PBS and incubated with both Alexa-488 anti-rat (1:250, 712-545-153, Jackson ImmunoResearch) and Rhodamine anti-mouse (1:250, 715-295-151, Jackson ImmunoResearch) secondary antibodies for 1 h at room temperature. All cell images were captured on a Zeiss Axio Imager M2 microscope with a Zeiss Axiocam 705 mono camera and processed using Zeiss ZEN lite software. DNA fiber analysis was carried out with ImageJ software (NIH).

### Immunofluorescence

Immunofluorescence (IF) was performed as described^77,78^. To detect nascent ssDNA, cells seeded on coverslips were incubated with 10 μM BrdU for 10 min prior to treatment with or without 4 mM HU for 4 h. To detect parental ssDNA, cells seeded on coverslips were incubated with 10 μM BrdU for 20 h and then released into fresh media for 2 h prior to treatment with or without 4 mM HU for 4 h. Subsequently, cells on coverslips were incubated with 0.5% Triton X-100 in PBS for 5 min at 4°C, washed with PBS, and then fixed with PBS-buffered 4% paraformaldehyde before proceeding with regular IF as described^77,78^. To detect EdU in interphase, cells seeded on coverslips were incubated with 10 μM EdU for 10 min prior to treatment with or without 4 mM HU. Subsequently, cells on coverslips were fixed, washed with PBS and then incubated with freshly prepared Click-iT reaction buffer (2 mM CuSO4, 10 μM biotin-PEG3-azide, 10 mM Ascorbic acid) for 10 min at room temperature. Coverslips were then washed in PBS twice, followed by regular IF as described^77,78^. To detect EdU in early mitosis, fixed prometaphase cells were treated with 0.5% Triton X-100 in PBS for 20 min, blocked with 0.5% bovine serum albumin (Sigma), and 0.2% gelatin (Sigma) for 30 min, and then incubated with freshly prepared Click-iT reaction buffer for 50 min at room temperature as described^79^. To detect HU-induced SMARCAL1 foci, cells were pre-extracted with cold CSK Buffer (10 mM PIPES pH 7.0, 100 mM NaCl, 300 mM Sucrose, 3 mM MgCl_2_, 0.5% Triton X-100) for 5 min prior to fixation as described^74^. All cell images were captured on a Zeiss Axio Imager M2 microscope with a Zeiss Axiocam 705 mono camera and processed using Zeiss ZEN lite software.

### Flow cytometer analyses

For GFP reporter assays, U2OS DR-GFP reporter cells were first transfected with control siRNA or siRNA against ASCC3. 24 h later, they were transfected with I-SceI and pCherry with a 9:1 ratio. 48 h post I-SceI transfection, cells were harvested, fixed, and subjected to flow cytometer analysis as described^58,77^. Cherry expression was used as a transfection efficiency control. A total of 20,000 events per cell line were scored for each independent experiment using a BD^TM^ Accuri C6 Plus Flow Cytometer.

For cell cycle analysis, following harvesting, U2OS parental and ASCC3-KO cells were washed twice with cold PBS and then fixed in 80% ethanol at 4°C for overnight. Fixed cells were incubated in PBS containing 100 μg/mL RNase A and 50 μg/mL propidium iodide at 37°C for 30 minutes, followed by flow cytometry analysis using a BD^TM^ Accuri C6 Plus Flow Cytometer. Flow cytometer data were exported and processed in FlowJo (v10.9).

### Immunoprecipitation and immunoblotting

Immunoprecipitation (IP) with endogenous ASCC2 and ASCC3 proteins was carried out as described^58,74^ with minor modifications. Briefly, HCT116 cells were lysed in NETN buffer [20 mM Tris-HCl, pH 8.0, 100 mM NaCl, 1 mM EDTA, 0.5% Nonidet^TM^ P-40 Substitute (Sigma), 1 mM PMSF, 1 μg/ml aprotinin, 1 μg/ml leupeptin, 1 μg/ml pepstatin, 1 mM NaF, 1 mM NaVO4, 50 mM Na-β-glycerolphosphate, 20 μM N-ethylmaleimide, 1 μM Iodoacetamine] on ice for 30 min. Following centrifugation at 13,000 rpm for 10 min at 4°C, the supernatant (cell lysate) was transferred to a clean tube. For each IP, 2 mg of cell lysate was first incubated with 1 μl of primary antibody overnight at 4 °C and then with protein G beads (30 μl) for 1 h at 4°C. Subsequently, precipitates were then washed 4 times in NETN buffer containing 300 mM NaCl and immunoblotted with indicated antibodies. Immunoprecipitation (IP) with exogenously expressed proteins were carried out as described^80^ with minor modifications. HEK293T cells grown on 6-cm plates with 70-80% confluency were transfected with a total of 8 μg DNA of the indicated constructs using Lipofectamine 2000 (Invitrogen) according to the manufacturer’s protocol. For co-transfection of epitope-tagged RPA32/RPA70 and epitope-tagged full length ASCC3, a DNA mixture of 1:3 was used. For co-transfection of Myc-ASCC3 deletion alleles and HA-RPA70, a ratio of 5:3 was used. For each IP, 5 μl of an anti-Myc antibody or 10 μl of an anti-HA antibody was used. To test for the interaction between HA-RPA70 and endogenous ASCC2/ASCC3 upon replication stress, 20 h post transfection, cells were treated with 4 mM HU for 4 h prior to preparation of cell lysates. For DNase I treatment, the cell lysate was incubated with 50 units of DNase I (EN0521, ThermoFisher) at 37°C for 20 min in the presence of 1x DNase I buffer (B43, ThermoFisher). Subsequently, anti-HA antibody was added, and the mixture was incubated in the presence of DNase I at 4°C overnight. Immunoblotting was performed as described^81^.

Immunoprecipitation of polyubiquitylated PCNA by HA-tagged ASCC2 was carried out as described^29^. Briefly, 2.5 million HEK293T cells were seeded per 10-cm plate. The following day, cells were transfected with 60 pmol siRNA against USP1. 24 post later, cells were transfected with 16 μg of either vector alone, pLPC-NFH2-ASCC2, or pLPC-NFH2-ASCC2 carrying triple L^478^L^479^P^480^-AAA mutations. 48 post initial transfection, cells were treated with 10 μM ATR inhibitor VE-821 for 30 min, gently washed with warm PBS, and exposed to 30 J/m² UV. Cells were then incubated in fresh media containing 10 µM ATRi for 4 h. Subsequently, cells were fixed in 0.25% formaldehyde in PBS for 15 min at room temperature and quenched with 0.125 M glycine for 5 min. Cells were harvested by scraping, washed twice with cold PBS, and then incubated in lysis buffer (50 mM Tris-HCl pH 8.0, 10 mM EDTA pH 8.0, 1% SDS, 1 mM PMSF, 1 μg/ml aprotinin, 1 μg/ml leupeptin, 1 μg/ml pepstatin) for 20 min on ice. Following two washes with cold TE buffer, cells were sonicated in TE buffer containing 0.1% SDS. Following centrifugation, the cell lysate was supplemented with 150 mM NaCl and 0.1% NP-40, followed by overnight incubation with 20 µL anti-HA antibody at 4°C.

### Molecular cloning of ASCC3^HR^- and TRIP4-expressing baculovirus constructs

The DNA sequence encoding the dual-cassette helicase region of ASCC3 (residues 401-2202; ASCC3^HR^ WT)^42^ was cloned into a pFL vector for expression under the control of the very late polyhedrin promoter. This promoter facilitated the expression of N-terminally His_10_-tagged, TEV-cleavable fusion proteins via recombinant baculoviruses in insect cells. Mutations resulting in the ASCC3^HR^ variant (D611A-D1453A) were introduced using the QuikChange II XL Site-Directed Mutagenesis Kit (Agilent). The expression cassettes were integrated into the MultiBac baculoviral genome through Tn7 transposition within a lacZα gene, enabling the selection of recombinants via blue/white screening.

A DNA fragment encoding full-length TRIP4 was PCR-amplified from a synthetic gene (IDT) and inserted into the pIDK vector (EMBL, Heidelberg) for expression as an N-terminally TwinStrep-tagged, TEV-cleavable fusion protein. For co-expression, the pIDK-trip4 construct was Cre-recombined with pFL-ASCC3^HR^ (WT or D611A-D1453A), generating a recombinant baculovirus for insect cell expression. All constructs were confirmed by Sanger sequencing to ensure accuracy.

### Recombinant protein expression and purification

ASCC3^HR^ (WT or D611A-D1453A) was expressed in High Five cells (Thermo Fisher Scientific B85502) via recombinant baculoviruses which were produced in Sf9 cells (Thermo Fisher Scientific 11496015)^42,43^. The cell pellets were resuspended in lysis buffer containing 500 mM NaCl, 20 mM HEPES-NaOH, pH 7.5, 10 mM imidazole, 1 mM DTT, 8.6% (v/v) glycerol (buffer A) supplemented with cOmplete^TM^ protease inhibitors (Roche) and lysed by sonication using a Sonopuls Ultrasonic Homogenizer HD (Bandelin). The lysate containing the soluble protein was centrifuged at 215,000 xg for 1 h and filtered through 0.8 µm pore size membrane filters (Millipore). The Ni^2+^-NTA resin was washed using buffer A in a gravity flow column, the resins were incubated with cleared lysate for 1 h and the protein of interest (POI) was eluted with buffer A containing 400 mM imidazole. The His_10_-tag was removed by incubating the POI with 1/10 (w/w) TEV protease overnight at 4°C and was simultaneously dialyzed against 20 mM HEPES-NaOH, pH 7.5, 500 mM NaCl, 1 mM DTT, 8.6% (v/v) glycerol (dialysis buffer). The sample was then diluted to 100 mM NaCl using 20 mM HEPES-NaOH, pH 7.5, 1 mM DTT, 8.6% (v/v) glycerol and loaded onto a HiTrap Heparin HP column (Cytiva), pre-equilibrated with buffer A containing 100 mM NaCl. After washing with buffer A containing 100 mM NaCl, the POI was eluted with a linear gradient to buffer A containing 1.5 M NaCl. The fractions containing the POI were pooled and concentrated with a centrifugal concentrator (100 kDa molecular mass cutoff). The concentrated sample was further purified by size-exclusion chromatography (SEC) on a Superdex 200 10/300 GL column (Cytiva) in 20 mM HEPES-NaOH, pH 7.5, 250 mM NaCl, 5 % (v/v) glycerol, 1 mM DTT (SEC buffer). Fractions containing the POI were combined, concentrated, aliquoted, flash-frozen in liquid nitrogen, and stored at −80 °C.

For the preparation of the ASCC3^HR^ (WT or D611A-D1453A)-TRIP4 complex, TRIP4 was co-produced with ASCC3^HR^ in High Five cells^43^. The cell pellets were re-suspended in lysis buffer containing 500 mM NaCl, 20 mM HEPES-NaOH, pH 7.5, 10 mM imidazole, 1 mM DTT, 8.6% (v/v) glycerol (buffer A) supplemented with cOmplete^TM^ protease inhibitors (Roche) and lysed by sonication using a Sonopuls Ultrasonic Homogenizer HD (Bandelin). The lysate containing the soluble protein was centrifuged at 215,000 xg for 1 h and filtered through 0.8 µm pore size membrane filters (Millipore). The Ni^2+^-NTA resin was washed using buffer A in a gravity flow column, the resins were incubated with cleared lysate for 1 h and the POI were eluted with buffer A containing 400 mM imidazole. Fractions enriched with POI were incubated with strep-Tactin resins for 1 h and eluted using buffer A containing 2.5 mM desthiobiotin. The fractions containing the POI were pooled and concentrated with a centrifugal concentrator (50 kDa molecular mass cutoff). The concentrated sample was further purified by SEC on a Superdex 200 10/300 GL column (Cytiva) in SEC buffer. Fractions containing the POI were combined, concentrated, aliquoted, flash-frozen in liquid nitrogen, and stored at −80 °C.

### In vitro unwinding assays

DNA unwinding activities were evaluated using fluorescence-based stopped-flow experiments on an SX-20MV spectrometer (Applied Photophysics). The DNA substrate contained a 12-base pair duplex region and a 31-nucleotide 3’-single-stranded (ss) overhang. The short strand was labeled with an Alexa Fluor 488 fluorophore at the 3’-end, the complementary strand was labeled with a black hole quencher (BHQ-1) at the 5’-end (5’-**CGGCTCGCGGCC**-3’[Alexa488]; [BHQ-1]5’-**GGCCGCGAGCCG**GAAATTTAATTATAAACCAGACCGTCTCCTC-3’; complementary regions in bold). Experiments were carried out in 40 mM HEPES-NaOH, pH 7.5, 80 mM NaCl, 0.5 mM MgCl_2_ at 30°C. ASCC3^HR^ (WT or D611A-D1453A)-TRIP4 complex (250 nM) was incubated with duplex DNA (50 nM) for 5 minutes at 30°C. 60 µl of the protein-DNA mixture were then rapidly mixed with 60 µl of 4 mM ATP/MgCl_2_. Alexa Fluor 488 fluorescence emission was monitored for 20 minutes, using a 495 nm cutoff filter (KV 495, Schott). Traces from three technical replicates were averaged, baseline-corrected by subtracting the fluorescence value observed immediately after mixing, and normalized to the baseline-corrected maximum fluorescence of the highest-amplitude trace of the experimental series. Data were fitted to a double exponential equation (fraction unwound = *A*_fast_*(1 – exp(–*k*_fast_ * *t*)) + *A*_slow_ * (1 – exp(– *k*_slow_ * *t*)); *A*_fast/slow_, unwinding amplitudes of the fast/slow phases; *k*_fast/slow_, unwinding rate constants of the fast/slow phases [s^−1^]; *t*, time [s])^82^ using Prism software (version 8.0; GraphPad). Amplitude-weighted unwinding rate constants were calculated as *k*_uaw_ = (*A*_fast_ * *k*_fast_^2^ + *A*_slow_ * *k*_slow_^2^) / (*A*_fast_ * *k*_fast_ + *A*_slow_ * *k*_slow_)^82^. For the ASCC3^HR^ WT-TRIP4 complex, *k*_uaw_ was evaluated as 0.05 s^-1^, consistent with our previous findings^43^.

### In vitro assays of RPA accumulation on ssDNA

RPA accumulation was monitored using biotinylated DNA with an extended 3’-ssDNA tail under three different conditions: control (no ASCC3), ASCC3^HR^ WT-TRIP4, and ASCC3^HR^-D611A-D1453A-TRIP4. Duplex DNA was made by annealing 100mer oligonucleotide (5’-GGGCGAATTGGGCCCGACGTCGCATGCTCCTCTAGACTCGAGGAATTCGGTACCCCGGGTTCGAAATCGATAAGCTTACAGTCTCCATTTAAAGGACAAG-3’) with a 75mer oligonucleotide (5’ GCTTATCGATTTCGAACCCGGGGTACCGAATTCCTCGAGTCTAGAGGAGCATGCGACGTCGGGCCCAATTCGCCC-3’) complementary to its 5’-sequence. The annealed DNA (100 nM) was incubated with ASCC3^HR^ (WT or D611A-D1453A)-TRIP4 complex (575 nM) for 45 min at 37°C in the reaction buffer (25 mM MOPS (morpholine-propanesulfonic acid) pH 7.0, 60 mM KCl, 0.2% Tween-20, 2 mM DTT, 2.5 mM ATP, 5 mM MgCl_2_). Following this initial incubation, 250 nM of replication protein A (RPA) was added to each reaction, and the samples were further incubated at 37°C for 30 minutes. After the second incubation, input fractions were taken from the reaction before adding streptavidin bead. Then, reactions including beads was incubated 15 min at 37°C. The reaction tubes were then placed in a magnetic holder for 2 min to allow streptavidin beads to capture biotin-labeled oligonucleotides present in the reaction mixture. After bead capture, protein loading buffer was added to each sample. The samples were subjected to SDS-PAGE on a 4-12% gradient gel, run at 200 V for 30 min. The separated proteins were subsequently transferred onto a nitrocellulose membrane at 100 V for 1 h at room temperature. The membrane was blocked and then incubated overnight with a primary monoclonal mouse antibody against RPA70 (Calbiochem, #NA13, 1:1000). The next day, the membrane was incubated with a secondary anti-mouse antibody (1:10000) for 45 min at room temperature. Detection was performed using Western Lightning ECL substrate, and the signal was visualized using the iBright imaging system. The intensity of each band of RPA was calculated with Fiji (ImageJ) software.

### In vitro strand exchange assays

Strand exchange assays were performed using a duplex DNA with an extended 3’-ssDNA tail. Duplex DNA was made by annealing 100mer oligonucleotide (5’-GGGCGAATTGGGCCCGACGTCGCATGCTCCTCTAGACTCGAGGAATTCGGTACCCCGGGTTCGAAATCGATAAGCTTACAGTCTCCATTTAAAGGACAAG-3’) with a 27mer oligonucleotide (3’-CCCGCTTAACCCGGGCTGCAGCGTACG-5’) complementary to its 5’-sequence. A 35-mer 5’-^32^P-end-labelled nucleotide (5’-GCTTATCGATTTCGAACCCGGGGTACCGAATTCCT-3’) forming a dsDNA duplex was prepared by annealing with its complement 35mer oligonucleotide (5’-AGGAATTCGGTACCCCGGGTTCGAAATCGATAAGC-3’). Strand exchange reactions were performed with 1µM of 3’-ssDNA tail incubated with RAD51 (270 nM) for 5 min at 37°C in strand exchange buffer (25 mM MOPS (morpholine-propanesulfonic acid) pH 7.0, 60 mM KCl, 0.2% Tween-20, 2 mM DTT, 2 mM ATP, 2 mM MgCl_2_). Then 5’-^32^P-end-labeled homologous 35 bp duplex (350 nM) was added and incubated for 15 min at 37°C. Subsequently, an increasing concentration (150 nM, 300 nM, and 600 nM) of ASCC3^HR^ WT-TRIP4 or ASCC3^HR^-D611A-D1453A-TRIP4 was added to the reaction. After 45 min incubation, the product was deproteinized with proteinase K for 15 min incubation at 37°C. Samples were analyzed by electrophoresis through 8% PAGE in TBE buffer, dried onto chromatography paper and visualized by autoradiography.

### Statistical Analysis

A Student’s two-tailed unpaired t-test was used to derive all *P* values except for where specified.

## DATA AVAILABILITY

All data used in this study are available within the article, Supplementary files, or available from the authors upon request.

## SUPPLEMENTARY DATA

accompanies the paper.

## ACKNOWLEDGEMENTS

We would like to thank Titia de Lange for phoenix cells, Ramanujan Hegde for plasmids pCMV6-MYKDDK-ASCC2, pCMV6-MYKDDK-ASCC2-mut1, and pcDNA3.1-ASCC3-3xFLAG, Nima Mosammaparast for TRIP4 expression constructs, and Sunbin Liu for plasmid pIDK-Twin-StrepII-TRIP4.

This work was supported by funding from Canadian Institutes of Health Research to J.Y.M. (PJT526991) and X.-D.Z. (PJT159793), and from the Deutsche Forschungsgemeinschaft to M.C.W. (SFB1565, project number 469281184). A.K.B. was supported by NIH R35 GM141805. J.-Y.M. holds a Tier 1 Canada Research Chair in DNA Repair and Cancer Therapeutics. A.A. was supported by a PhD scholarship of the German Academic Exchange Service and S.V.S. was supported by a Cancer Research Center Jacques-Landry PhD Award.

## AUTHOR CONTRIBUTIONS

Conceptualization, J.R.W. and X.D.Z.; Methodology, A.K.B., M.C.W., J.Y.M, and X.D.Z.; Investigation, S.C., N.L.B., Y.C., A.A., J.R.W., and S.V.S; Writing – Original Draft, J.R.W. and X.D.Z.; Writing – Review & Editing, S.C., N.L.B, J.R.W., A.K.B., M.C.W., J.Y.M., and X.D.Z; Funding Acquisition, A.K.B., M.C.W., J.Y.M., X.D.Z.; Resources: A.K.B.; Supervision, M.C.W., J.Y.M, and X.D.Z.

## REFERENCES

1. Zeman, M.K., and Cimprich, K.A. (2014). Causes and consequences of replication stress. Nat Cell Biol 16, 2–9. 10.1038/ncb2897.

2. Gaillard, H., Garcia-Muse, T., and Aguilera, A. (2015). Replication stress and cancer. Nat Rev Cancer 15, 276–289. 10.1038/nrc3916.

3. da Costa, A., Chowdhury, D., Shapiro, G.I., D’Andrea, A.D., and Konstantinopoulos, P.A. (2023). Targeting replication stress in cancer therapy. Nat Rev Drug Discov 22, 38–58. 10.1038/s41573-022-00558-5.

4. Cybulla, E., and Vindigni, A. (2023). Leveraging the replication stress response to optimize cancer therapy. Nat Rev Cancer 23, 6–24. 10.1038/s41568-022-00518-6.

5. Saxena, S., and Zou, L. (2022). Hallmarks of DNA replication stress. Mol Cell 82, 2298–2314. 10.1016/j.molcel.2022.05.004.

6. Berti, M., Cortez, D., and Lopes, M. (2020). The plasticity of DNA replication forks in response to clinically relevant genotoxic stress. Nat Rev Mol Cell Biol 21, 633–651. 10.1038/s41580-020-0257-5.

7. Rickman, K., and Smogorzewska, A. (2019). Advances in understanding DNA processing and protection at stalled replication forks. J Cell Biol 218, 1096–1107. 10.1083/jcb.201809012.

8. Cimprich, K.A., and Cortez, D. (2008). ATR: an essential regulator of genome integrity. Nat Rev Mol Cell Biol 9, 616–627. 10.1038/nrm2450.

9. Quinet, A., Lemacon, D., and Vindigni, A. (2017). Replication Fork Reversal: Players and Guardians. Mol Cell 68, 830–833. 10.1016/j.molcel.2017.11.022.

10. Adar, S., Izhar, L., Hendel, A., Geacintov, N., and Livneh, Z. (2009). Repair of gaps opposite lesions by homologous recombination in mammalian cells. Nucleic Acids Res 37, 5737–5748. 10.1093/nar/gkp632.

11. Tirman, S., Quinet, A., Wood, M., Meroni, A., Cybulla, E., Jackson, J., Pegoraro, S., Simoneau, A., Zou, L., and Vindigni, A. (2021). Temporally distinct post-replicative repair mechanisms fill PRIMPOL-dependent ssDNA gaps in human cells. Mol Cell 81, 4026–4040 e4028. 10.1016/j.molcel.2021.09.013.

12. Edmunds, C.E., Simpson, L.J., and Sale, J.E. (2008). PCNA ubiquitination and REV1 define temporally distinct mechanisms for controlling translesion synthesis in the avian cell line DT40. Mol Cell 30, 519–529. 10.1016/j.molcel.2008.03.024.

13. Quinet, A., Tirman, S., Jackson, J., Svikovic, S., Lemacon, D., Carvajal-Maldonado, D., Gonzalez-Acosta, D., Vessoni, A.T., Cybulla, E., Wood, M., et al. (2020). PRIMPOL-Mediated Adaptive Response Suppresses Replication Fork Reversal in BRCA-Deficient Cells. Mol Cell 77, 461–474 e469. 10.1016/j.molcel.2019.10.008.

14. Vallerga, M.B., Mansilla, S.F., Federico, M.B., Bertolin, A.P., and Gottifredi, V. (2015). Rad51 recombinase prevents Mre11 nuclease-dependent degradation and excessive PrimPol-mediated elongation of nascent DNA after UV irradiation. Proc Natl Acad Sci U S A 112, E6624–6633. 10.1073/pnas.1508543112.

15. Zellweger, R., Dalcher, D., Mutreja, K., Berti, M., Schmid, J.A., Herrandor, R., Vindigni, A., and Lopes, M. (2015). Rad51-mediated replication fork reversal is a global response to genotoxic treatments in human cells. J Cell Biol 208, 563–579. 10.1083/jcb.201406099.

16. Berti, M., and Vindigni, A. (2016). Replication stress: getting back on track. Nat Struct Mol Biol 23, 103–109. 10.1038/nsmb.3163.

17. Kavlashvili, T., Liu, W., Mohamed, T.M., Cortez, D., and Dewar, J.M. (2023). Replication fork uncoupling causes nascent strand degradation and fork reversal. Nat Struct Mol Biol 30, 115–124. 10.1038/s41594-022-00871-y.

18. Adolph, M.B., and Cortez, D. (2024). Mechanisms and regulation of replication fork reversal. DNA Repair (Amst) 141, 103731. 10.1016/j.dnarep.2024.103731.

19. Machwe, A., Xiao, L., Groden, J., and Orren, D.K. (2006). The Werner and Bloom syndrome proteins catalyze regression of a model replication fork. Biochemistry 45, 13939–13946. 10.1021/bi0615487.

20. Chen, J., Wu, M., Yang, Y., Ruan, C., Luo, Y., Song, L., Wu, T., Huang, J., Yang, B., and Liu, T. (2024). TFIP11 promotes replication fork reversal to preserve genome stability. Nat Commun 15, 1262. 10.1038/s41467-024-45684-3.

21. Ralf, C., Hickson, I.D., and Wu, L. (2006). The Bloom’s syndrome helicase can promote the regression of a model replication fork. J Biol Chem 281, 22839–22846. 10.1074/jbc.M604268200.

22. Fugger, K., Mistrik, M., Neelsen, K.J., Yao, Q., Zellweger, R., Kousholt, A.N., Haahr, P., Chu, W.K., Bartek, J., Lopes, M., et al. (2015). FBH1 Catalyzes Regression of Stalled Replication Forks. Cell Rep 10, 1749–1757. 10.1016/j.celrep.2015.02.028.

23. Leung, W., Baxley, R.M., Moldovan, G.L., and Bielinsky, A.K. (2018). Mechanisms of DNA Damage Tolerance: Post-Translational Regulation of PCNA. Genes (Basel) 10. 10.3390/genes10010010.

24. Hoege, C., Pfander, B., Moldovan, G.L., Pyrowolakis, G., and Jentsch, S. (2002). RAD6-dependent DNA repair is linked to modification of PCNA by ubiquitin and SUMO. Nature 419, 135–141. 10.1038/nature00991.

25. Motegi, A., Liaw, H.J., Lee, K.Y., Roest, H.P., Maas, A., Wu, X., Moinova, H., Markowitz, S.D., Ding, H., Hoeijmakers, J.H., and Myung, K. (2008). Polyubiquitylation of proliferating cell nuclear antigen by HLTF and SHPRH prevents genomic instability from stalled replication forks. Proc Natl Acad Sci U S A 105, 12411–12416. 10.1073/pnas.0805685105.

26. Motegi, A., Sood, R., Moinova, H., Markowitz, S.D., Liu, P.P., and Myung, K. (2006). Human SHPRH suppresses genomic instability through proliferating cell nuclear antigen polyubiquitination. J Cell Biol 175, 703–708. 10.1083/jcb.200606145.

27. Yates, M., and Maréchal, A. (2018). Ubiquitylation at the Fork: Making and Breaking Chains to Complete DNA Replication. Int J Mol Sci 19, 2909. 10.3390/ijms19102909.

28. Vujanovic, M., Krietsch, J., Raso, M.C., Terraneo, N., Zellweger, R., Schmid, J.A., Taglialatela, A., Huang, J.W., Holland, C.L., Zwicky, K., et al. (2017). Replication Fork Slowing and Reversal upon DNA Damage Require PCNA Polyubiquitination and ZRANB3 DNA Translocase Activity. Mol Cell 67, 882–890 e885. 10.1016/j.molcel.2017.08.010.

29. Ciccia, A., Nimonkar, A.V., Hu, Y., Hajdu, I., Achar, Y.J., Izhar, L., Petit, S.A., Adamson, B., Yoon, J.C., Kowalczykowski, S.C., et al. (2012). Polyubiquitinated PCNA recruits the ZRANB3 translocase to maintain genomic integrity after replication stress. Mol Cell 47, 396–409. 10.1016/j.molcel.2012.05.024.

30. Jung, D.J., Sung, H.S., Goo, Y.W., Lee, H.M., Park, O.K., Jung, S.Y., Lim, J., Kim, H.J., Lee, S.K., Kim, T.S., et al. (2002). Novel transcription coactivator complex containing activating signal cointegrator 1. Mol Cell Biol 22, 5203–5211. 10.1128/mcb.22.14.5203-5211.2002.

31. Dango, S., Mosammaparast, N., Sowa, M.E., Xiong, L.J., Wu, F., Park, K., Rubin, M., Gygi, S., Harper, J.W., and Shi, Y. (2011). DNA unwinding by ASCC3 helicase is coupled to ALKBH3-dependent DNA alkylation repair and cancer cell proliferation. Mol Cell 44, 373–384. 10.1016/j.molcel.2011.08.039.

32. Brickner, J.R., Soll, J.M., Lombardi, P.M., Vågbø, C.B., Mudge, M.C., Oyeniran, C., Rabe, R., Jackson, J., Sullender, M.E., Blazosky, E., et al. (2017). A ubiquitin-dependent signalling axis specific for ALKBH-mediated DNA dealkylation repair. Nature 551, 389–393. 10.1038/nature24484.

33. Juszkiewicz, S., Speldewinde, S.H., Wan, L., Svejstrup, J.Q., and Hegde, R.S. (2020). The ASC-1 Complex Disassembles Collided Ribosomes. Mol Cell 79, 603–614.e608. 10.1016/j.molcel.2020.06.006.

34. Stoneley, M., Harvey, R.F., Mulroney, T.E., Mordue, R., Jukes-Jones, R., Cain, K., Lilley, K.S., Sawarkar, R., and Willis, A.E. (2022). Unresolved stalled ribosome complexes restrict cell-cycle progression after genotoxic stress. Mol Cell 82, 1557–1572.e1557. 10.1016/j.molcel.2022.01.019.

35. Chi, B., O’Connell, J.D., Iocolano, A.D., Coady, J.A., Yu, Y., Gangopadhyay, J., Gygi, S.P., and Reed, R. (2018). The neurodegenerative diseases ALS and SMA are linked at the molecular level via the ASC-1 complex. Nucleic Acid Res 46, 11939–11951. 10.1093/nar/gky1093.

36. Munoz, S., Blanco-Romero, E., Gonzalez-Acosta, D., Rodriguez-Acebes, S., Megias, D., Lopes, M., and Mendez, J. (2024). RAD51 restricts DNA over-replication from re-activated origins. EMBO J 43, 1043–1064. 10.1038/s44318-024-00038-z.

37. Nakamura, K., Kustatscher, G., Alabert, C., Hodl, M., Forne, I., Volker-Albert, M., Satpathy, S., Beyer, T.E., Mailand, N., Choudhary, C., et al. (2021). Proteome dynamics at broken replication forks reveal a distinct ATM-directed repair response suppressing DNA double-strand break ubiquitination. Mol Cell 81, 1084–1099 e1086. 10.1016/j.molcel.2020.12.025.

38. Alabert, C., Bukowski-Wills, J.C., Lee, S.B., Kustatscher, G., Nakamura, K., de Lima Alves, F., Menard, P., Mejlvang, J., Rappsilber, J., and Groth, A. (2014). Nascent chromatin capture proteomics determines chromatin dynamics during DNA replication and identifies unknown fork components. Nat Cell Biol 16, 281–293. 10.1038/ncb2918.

39. Wessel, S.R., Mohni, K.N., Luzwick, J.W., Dungrawala, H., and Cortez, D. (2019). Functional Analysis of the Replication Fork Proteome Identifies BET Proteins as PCNA Regulators. Cell Rep 28, 3497–3509 e3494. 10.1016/j.celrep.2019.08.051.

40. Srivastava, M., Chen, Z., Zhang, H., Tang, M., Wang, C., Jung, S.Y., and Chen, J. (2018). Replisome Dynamics and Their Functional Relevance upon DNA Damage through the PCNA Interactome. Cell Rep 25, 3869–3883 e3864. 10.1016/j.celrep.2018.11.099.

41. Soll, J.M., Brickner, J.R., Mudge, M.C., and Mosammaparast, N. (2018). RNA ligase-like domain in activating signal cointegrator 1 complex subunit 1 (ASCC1) regulates ASCC complex function during alkylation damage. J Biol Chem 293, 13524–13533. 10.1074/jbc.RA117.000114.

42. Jia, J., Absmeier, E., Holton, N., Pietrzyk-Brzezinska, A.J., Hackert, P., Bohnsack, K.E., Bohnsack, M.T., and Wahl, M.C. (2020). The interaction of DNA repair factors ASCC2 and ASCC3 is affected by somatic cancer mutations. Nat Commun 11, 5535. 10.1038/s41467-020-19221-x.

43. Jia, J., Hilal, T., Bohnsack, K.E., Chernev, A., Tsao, N., Bethmann, J., Arumugam, A., Parmely, L., Holton, N., Loll, B., et al. (2023). Extended DNA threading through a dual-engine motor module of the activating signal co-integrator 1 complex. Nat Commun 14, 1886. 10.1038/s41467-023-37528-3.

44. Motegi, A., Sood, R., Moinova, H., Markowitz, S.D., Liu, P.P., and Myung, K. (2006). Human SHPRH suppresses genomic instability through proliferating cell nuclear antigen polyubiquitination. J Cell Biol 175, 703–708. 10.1083/jcb.200606145.

45. Leung, W., Baxley, R.M., Traband, E., Chang, Y.C., Rogers, C.B., Wang, L., Durrett, W., Bromley, K.S., Fiedorowicz, L., Thakar, T., et al. (2023). FANCD2-dependent mitotic DNA synthesis relies on PCNA K164 ubiquitination. Cell Rep 42, 113523. 10.1016/j.celrep.2023.113523.

46. Moore, C.E., Yalcindag, S.E., Czeladko, H., Ravindranathan, R., Wijesekara Hanthi, Y., Levy, J.C., Sannino, V., Schindler, D., Ciccia, A., Costanzo, V., and Elia, A.E.H. (2023). RFWD3 promotes ZRANB3 recruitment to regulate the remodeling of stalled replication forks. J Cell Biol 222. 10.1083/jcb.202106022.

47. Brun, J., Chiu, R.K., Wouters, B.G., and Gray, D.A. (2010). Regulation of PCNA polyubiquitination in human cells. BMC Res Notes 3, 85. 10.1186/1756-0500-3-85.

48. Huang, T.T., Nijman, S.M., Mirchandani, K.D., Galardy, P.J., Cohn, M.A., Haas, W., Gygi, S.P., Ploegh, H.L., Bernards, R., and D’Andrea, A.D. (2006). Regulation of monoubiquitinated PCNA by DUB autocleavage. Nat Cell Biol 8, 339–347. 10.1038/ncb1378.

49. Yang, X.H., and Zou, L. (2009). Dual functions of DNA replication forks in checkpoint signaling and PCNA ubiquitination. Cell Cycle 8, 191–194. 10.4161/cc.8.2.7357.

50. Ciccia, A., Bredemeyer, A.L., Sowa, M.E., Terret, M.E., Jallepalli, P.V., Harper, J.W., and Elledge, S.J. (2009). The SIOD disorder protein SMARCAL1 is an RPA-interacting protein involved in replication fork restart. Genes Dev 23, 2415–2425. 10.1101/gad.1832309.

51. Bétous, R., Mason, A.C., Rambo, R.P., Bansbach, C.E., Badu-Nkansah, A., Sirbu, B.M., Eichman, B.F., and Cortez, D. (2012). SMARCAL1 catalyzes fork regression and Holliday junction migration to maintain genome stability during DNA replication. Genes Dev 26, 151–162. 10.1101/gad.178459.111.

52. Yusufzai, T., Kong, X., Yokomori, K., and Kadonaga, J.T. (2009). The annealing helicase HARP is recruited to DNA repair sites via an interaction with RPA. Genes Dev 23, 2400–2404. 10.1101/gad.1831509.

53. Taglialatela, A., Alvarez, S., Leuzzi, G., Sannino, V., Ranjha, L., Huang, J.W., Madubata, C., Anand, R., Levy, B., Rabadan, R., et al. (2017). Restoration of Replication Fork Stability in BRCA1- and BRCA2-Deficient Cells by Inactivation of SNF2-Family Fork Remodelers. Mol Cell 68, 414–430 e418. 10.1016/j.molcel.2017.09.036.

54. Kolinjivadi, A.M., Sannino, V., De Antoni, A., Zadorozhny, K., Kilkenny, M., Techer, H., Baldi, G., Shen, R., Ciccia, A., Pellegrini, L., et al. (2017). Smarcal1-Mediated Fork Reversal Triggers Mre11-Dependent Degradation of Nascent DNA in the Absence of Brca2 and Stable Rad51 Nucleofilaments. Mol Cell 67, 867–881 e867. 10.1016/j.molcel.2017.07.001.

55. Mijic, S., Zellweger, R., Chappidi, N., Berti, M., Jacobs, K., Mutreja, K., Ursich, S., Ray Chaudhuri, A., Nussenzweig, A., Janscak, P., and Lopes, M. (2017). Replication fork reversal triggers fork degradation in BRCA2-defective cells. Nat Commun 8, 859. 10.1038/s41467-017-01164-5.

56. Taglialatela, A., Leuzzi, G., Sannino, V., Cuella-Martin, R., Huang, J.W., Wu-Baer, F., Baer, R., Costanzo, V., and Ciccia, A. (2021). REV1-Polzeta maintains the viability of homologous recombination-deficient cancer cells through mutagenic repair of PRIMPOL-dependent ssDNA gaps. Mol Cell 81, 4008–4025 e4007. 10.1016/j.molcel.2021.08.016.

57. Piberger, A.L., Bowry, A., Kelly, R.D.W., Walker, A.K., Gonzalez-Acosta, D., Bailey, L.J., Doherty, A.J., Mendez, J., Morris, J.R., Bryant, H.E., and Petermann, E. (2020). PrimPol-dependent single-stranded gap formation mediates homologous recombination at bulky DNA adducts. Nat Commun 11, 5863. 10.1038/s41467-020-19570-7.

58. Batenburg, N.L., Walker, J.R., Noordermeer, S.M., Moatti, N., Durocher, D., and Zhu, X.D. (2017). ATM and CDK2 control chromatin remodeler CSB to inhibit RIF1 in DSB repair pathway choice. Nat Commun 8, 1921. 10.1038/s41467-017-02114-x.

59. Escribano-Diaz, C., Orthwein, A., Fradet-Turcotte, A., Xing, M., Young, J.T., Tkac, J., Cook, M.A., Rosebrock, A.P., Munro, M., Canny, M.D., et al. (2013). A cell cycle-dependent regulatory circuit composed of 53BP1-RIF1 and BRCA1-CtIP controls DNA repair pathway choice. Mol Cell 49, 872–883. 10.1016/j.molcel.2013.01.001.

60. Tonzi, P., Yin, Y., Lee, C.W.T., Rothenberg, E., and Huang, T.T. (2018). Translesion polymerase kappa-dependent DNA synthesis underlies replication fork recovery. Elife 7. 10.7554/eLife.41426.

61. Bhowmick, R., Minocherhomji, S., and Hickson, I.D. (2016). RAD52 Facilitates Mitotic DNA Synthesis Following Replication Stress. Mol Cell 64, 1117–1126. 10.1016/j.molcel.2016.10.037.

62. Mocanu, C., Karanika, E., Fernández-Casañas, M., Herbert, A., Olukoga, T., Özgürses, M.E., and Chan, K.L. (2022). DNA replication is highly resilient and persistent under the challenge of mild replication stress. Cell Rep 39, 110701. 10.1016/j.celrep.2022.110701.

63. Nandi, B., Talluri, S., Kumar, S., Yenumula, C., Gold, J.S., Prabhala, R., Munshi, N.C., and Shammas, M.A. (2019). The roles of homologous recombination and the immune system in the genomic evolution of cancer. J Transl Sci 5. 10.15761/JTS.1000282.

64. Aguilera, A., and Gomez-Gonzalez, B. (2008). Genome instability: a mechanistic view of its causes and consequences. Nat Rev Genet 9, 204–217. 10.1038/nrg2268.

65. Matos-Rodrigues, G., Guirouilh-Barbat, J., Martini, E., and Lopez, B.S. (2021). Homologous recombination, cancer and the ‘RAD51 paradox’. NAR Cancer 3, zcab016. 10.1093/narcan/zcab016.

66. Fugger, K., Mistrik, M., Danielsen, J.R., Dinant, C., Falck, J., Bartek, J., Lukas, J., and Mailand, N. (2009). Human Fbh1 helicase contributes to genome maintenance via pro- and anti-recombinase activities. J Cell Biol 186, 655–663. 10.1083/jcb.200812138.

67. Dixit, S., Nagraj, T., Bhattacharya, D., Saxena, S., Sahoo, S., Chittela, R.K., Somyajit, K., and Nagaraju, G. (2024). RTEL1 helicase counteracts RAD51-mediated homologous recombination and fork reversal to safeguard replicating genomes. Cell Rep 43, 114594. 10.1016/j.celrep.2024.114594.

68. Bugreev, D.V., Yu, X., Egelman, E.H., and Mazin, A.V. (2007). Novel pro- and anti-recombination activities of the Bloom’s syndrome helicase. Genes Dev 21, 3085–3094. 10.1101/gad.1609007.

69. Mason, J.M., Chan, Y.L., Weichselbaum, R.W., and Bishop, D.K. (2019). Non-enzymatic roles of human RAD51 at stalled replication forks. Nat Commun 10, 4410. 10.1038/s41467-019-12297-0.

70. Hashimoto, Y., Ray Chaudhuri, A., Lopes, M., and Costanzo, V. (2010). Rad51 protects nascent DNA from Mre11-dependent degradation and promotes continuous DNA synthesis. Nat Struct Mol Biol 17, 1305–1311. 10.1038/nsmb.1927.

71. Schlacher, K., Christ, N., Siaud, N., Egashira, A., Wu, H., and Jasin, M. (2011). Double-strand break repair-independent role for BRCA2 in blocking stalled replication fork degradation by MRE11. Cell 145, 529–542. 10.1016/j.cell.2011.03.041.

72. Chappidi, N., Nascakova, Z., Boleslavska, B., Zellweger, R., Isik, E., Andrs, M., Menon, S., Dobrovolna, J., Balbo Pogliano, C., Matos, J., et al. (2020). Fork Cleavage-Religation Cycle and Active Transcription Mediate Replication Restart after Fork Stalling at Co-transcriptional R-Loops. Mol Cell 77, 528–541 e528. 10.1016/j.molcel.2019.10.026.

73. Berti, M., Ray Chaudhuri, A., Thangavel, S., Gomathinayagam, S., Kenig, S., Vujanovic, M., Odreman, F., Glatter, T., Graziano, S., Mendoza-Maldonado, R., et al. (2013). Human RECQ1 promotes restart of replication forks reversed by DNA topoisomerase I inhibition. Nat Struct Mol Biol 20, 347–354. 10.1038/nsmb.2501.

74. Batenburg, N.L., Sowa, D.J., Walker, J.R., Andres, S.N., and Zhu, X.D. (2024). CSB and SMARCAL1 compete for RPA32 at stalled forks and differentially control the fate of stalled forks in BRCA2-deficient cells. Nucleic Acids Res 52, 5067–5087. 10.1093/nar/gkae154.

75. Batenburg, N.L., Mersaoui, S.Y., Walker, J.R., Coulombe, Y., Hammond-Martel, I., Wurtele, H., Masson, J.Y., and Zhu, X.D. (2021). Cockayne syndrome group B protein regulates fork restart, fork progression and MRE11-dependent fork degradation in BRCA1/2-deficient cells. Nucleic Acids Res 49, 12836–12854. 10.1093/nar/gkab1173.

76. Batenburg, N.L., Walker, J.R., and Zhu, X.D. (2023). CSB Regulates Pathway Choice in Response to DNA Replication Stress Induced by Camptothecin. Int J Mol Sci 24. 10.3390/ijms241512419.

77. Batenburg, N.L., Thompson, E.L., Hendrickson, E.A., and Zhu, X.D. (2015). Cockayne syndrome group B protein regulates DNA double-strand break repair and checkpoint activation. EMBO J. 34, 1399–1416. 10.15252/embj.201490041.

78. Batenburg, N.L., Mitchell, T.R., Leach, D.M., Rainbow, A.J., and Zhu, X.D. (2012). Cockayne Syndrome group B protein interacts with TRF2 and regulates telomere length and stability. Nucleic Acids Res. 40, 9661–9674. 10.1093/nar/gks745.

79. Cui, S., Walker, J.R., Batenburg, N.L., and Zhu, X.D. (2022). Cockayne syndrome group B protein uses its DNA translocase activity to promote mitotic DNA synthesis. DNA Repair (Amst) 116, 103354. 10.1016/j.dnarep.2022.103354.

80. Batenburg, N.L., Mitchell, T.R., Leach, D.M., Rainbow, A.J., and Zhu, X.D. (2012). Cockayne Syndrome group B protein interacts with TRF2 and regulates telomere length and stability. Nucleic Acids Res 40, 9661–9674. 10.1093/nar/gks745.

81. Batenburg, N.L., Walker, J.R., Coulombe, Y., Sherker, A., Masson, J.Y., and Zhu, X.D. (2019). CSB interacts with BRCA1 in late S/G2 to promote MRN- and CtIP-mediated DNA end resection. Nucleic Acids Res 47, 10678–10692. 10.1093/nar/gkz784.

82. Ozes, A.R., Feoktistova, K., Avanzino, B.C., Baldwin, E.P., and Fraser, C.S. (2014). Real-time fluorescence assays to monitor duplex unwinding and ATPase activities of helicases. Nat Protoc 9, 1645–1661. 10.1038/nprot.2014.112.

