## Supplementary Figures for "The Ski2 helicase ASCC3 unwinds DNA upon fork stalling to control replication stress responses"

**SUPPLEMENTARY FIGURES S1-S9**

**Supplementary Figure S1 Cui et al.**

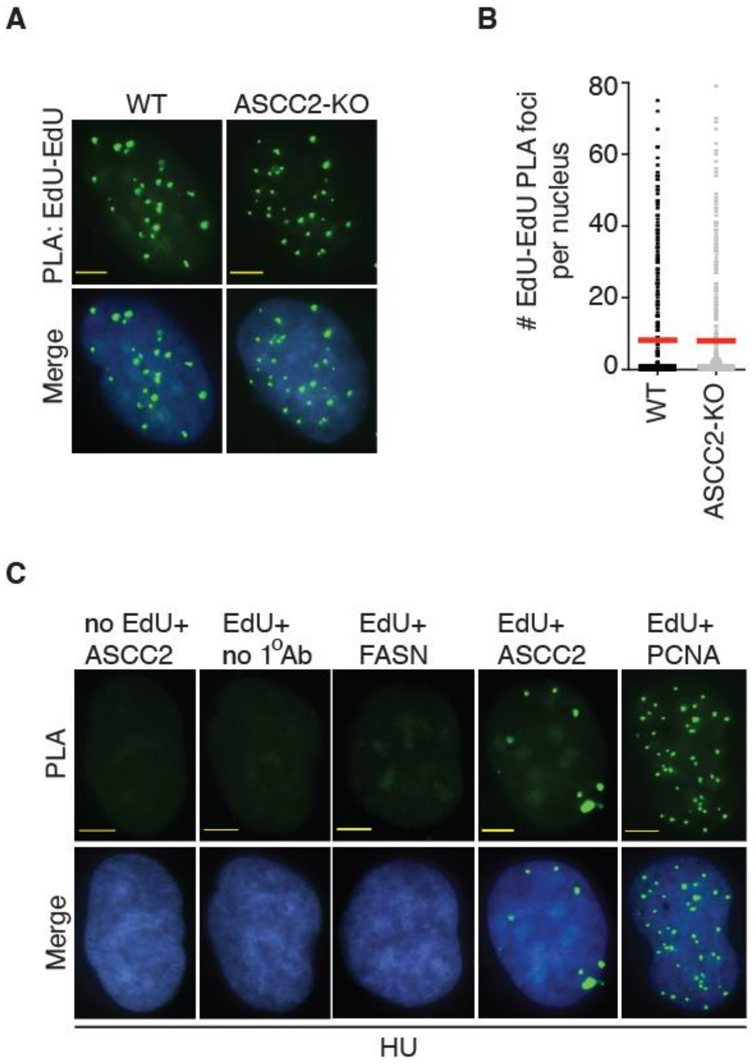

**Figure S1**. Recruitment of ASCC2 to stalled forks is not due to non-specific effects. (**A**) Representative images of EdU-EdU PLA foci formation in U2OS WT and ASCC2-KO. (**B**) Quantification of EdU-EdU PLA foci formation from (A). This PLA experiment was performed once. A total of 481-496 cells per condition were analyzed. (**C**) Representative images of PLA foci formation in U2OS treated with HU. PLA assays were performed in several conditions as indicated above the images.

**Supplementary Figure S2 Cui et al.**

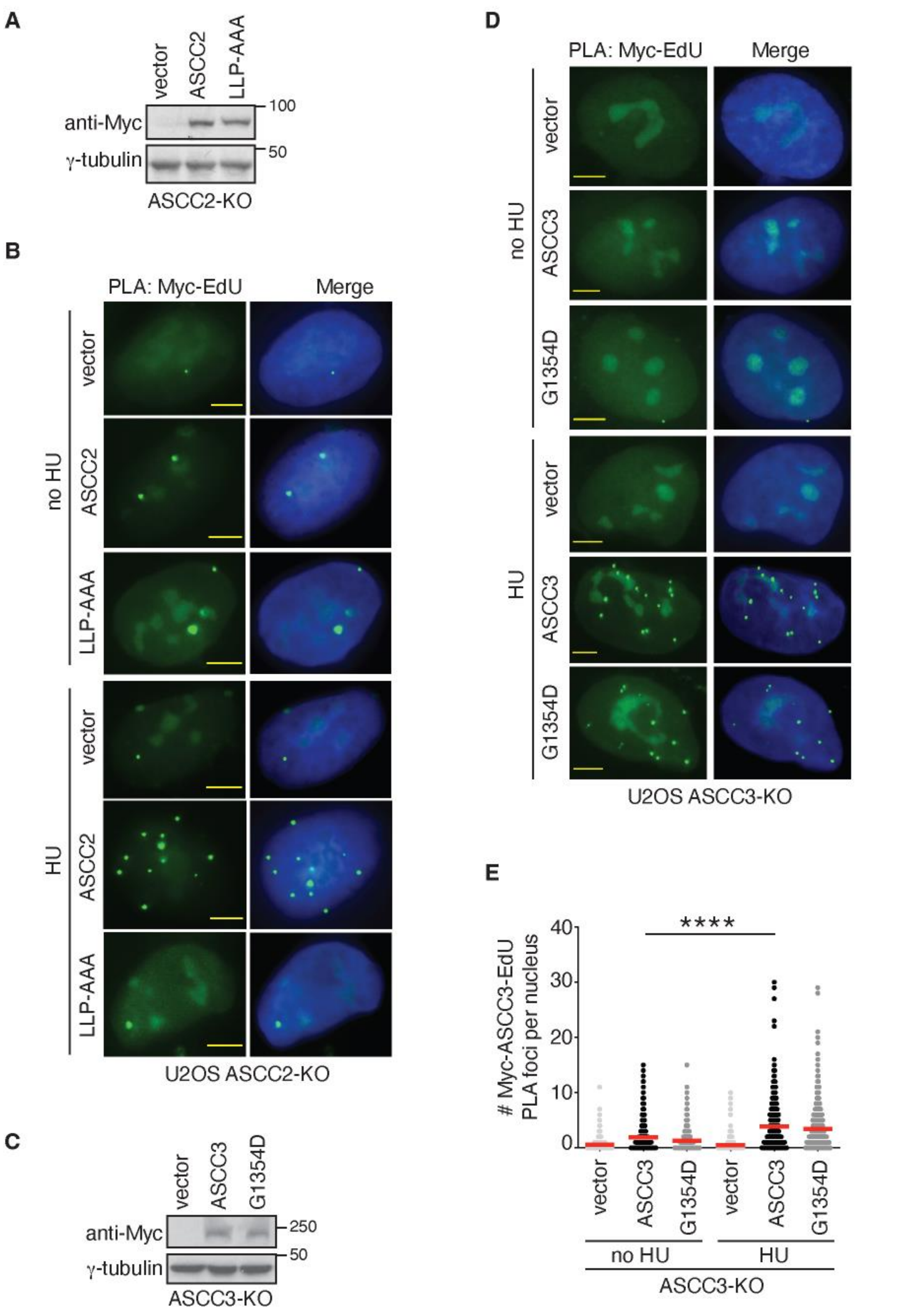

**Figure S2**. ASCC2 and ASCC3 are recruited to stalled forks, the former requiring its ubiquitin binding activity. (**A**) Western blot analyses of U2OS ASCC2-KO cells expressing various Myc-ASCC2 alleles as indicated. Immunoblotting was performed with anti-Myc and anti-γ-tubulin antibodies. (**B**) Representative images of Myc-EdU PLA foci formation in no HU- or HU-treated U2OS ASCC2-KO cells expressing various Myc-ASCC2 alleles as indicated. (**C**) Western blot analyses of U2OS ASCC3-KO cells expressing various Myc-ASCC3 alleles as indicated. Immunoblotting was performed with anti-Myc and anti-γ-tubulin antibodies. (**D**) Representative images of Myc-EdU PLA foci formation in no HU- or HU-treated U2OS ASCC3-KO cells expressing various Myc-ASCC3 alleles as indicated. (**E**) Quantification of Myc-ASCC3-EdU PLA foci formation in no HU- or HU-treated U2OS ASCC3-KO cells expressing various Myc-ASCC3 alleles as indicated. This PLA experiment was performed once. Data are represented as scatter plot graphs with the mean indicated. A total of 801-813 cells per condition were analyzed. The *P*-value was determined using a non-parametric Mann-Whitney rank-sum *t*-test. ****$P<0.0001$.

**Supplementary Figure S3 Cui et al.**

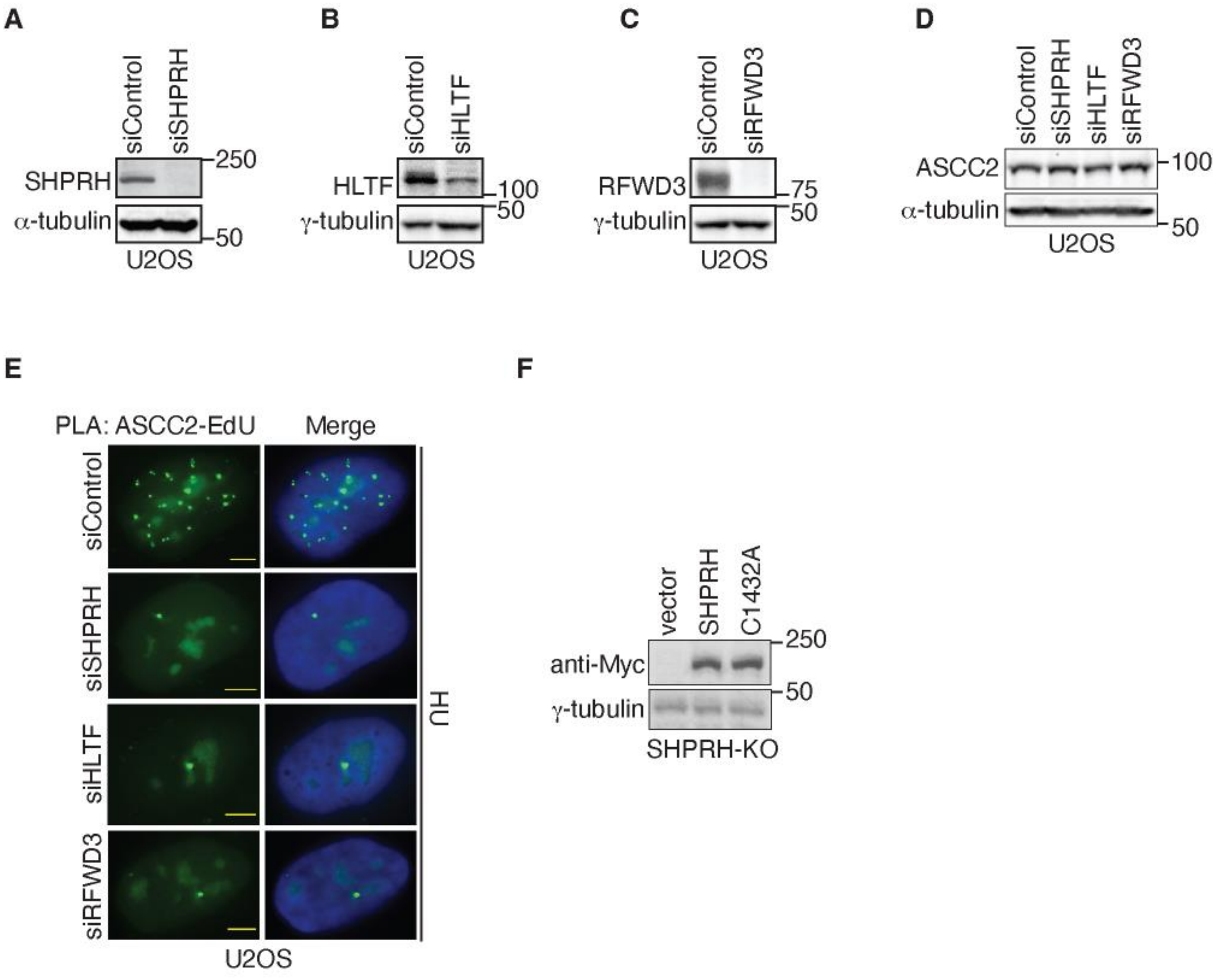

**Figure S3**. SHPRH, HLTF, and RFWD3 mediate recruitment of ASCC2 to stalled forks. (**A**) Western blot analyses of U2OS cells transfected with siControl or siSHPRH. Immunoblotting was performed with anti-SHPRH and anti-γ-tubulin antibodies. (**B**) Western blot analyses of U2OS cells transfected with siControl or siHLTF. Immunoblotting was performed with anti-HLTF and anti-γ-tubulin antibodies. (**C**) Western blot analyses of U2OS cells transfected with siControl or siRFWD3. Immunoblotting was performed with anti-RFWD3 and anti-γ-tubulin antibodies. (**D**) Western blot analyses of U2OS cells transfected with indicated siRNAs. Immunoblotting was performed with anti-ASCC2 and anti-α-tubulin antibodies. (**E**) Representative images of ASCC2-EdU PLA foci formation in HU-treated U2OS expressing indicated siRNAs. (**F**) Western blot analyses of U2OS SHPRH-KO cells expressing various Myc-SHPRH alleles as indicated. Immunoblotting was performed with anti-Myc and anti-γ-tubulin antibodies.

**Supplementary Figure S4 Cui et al.**

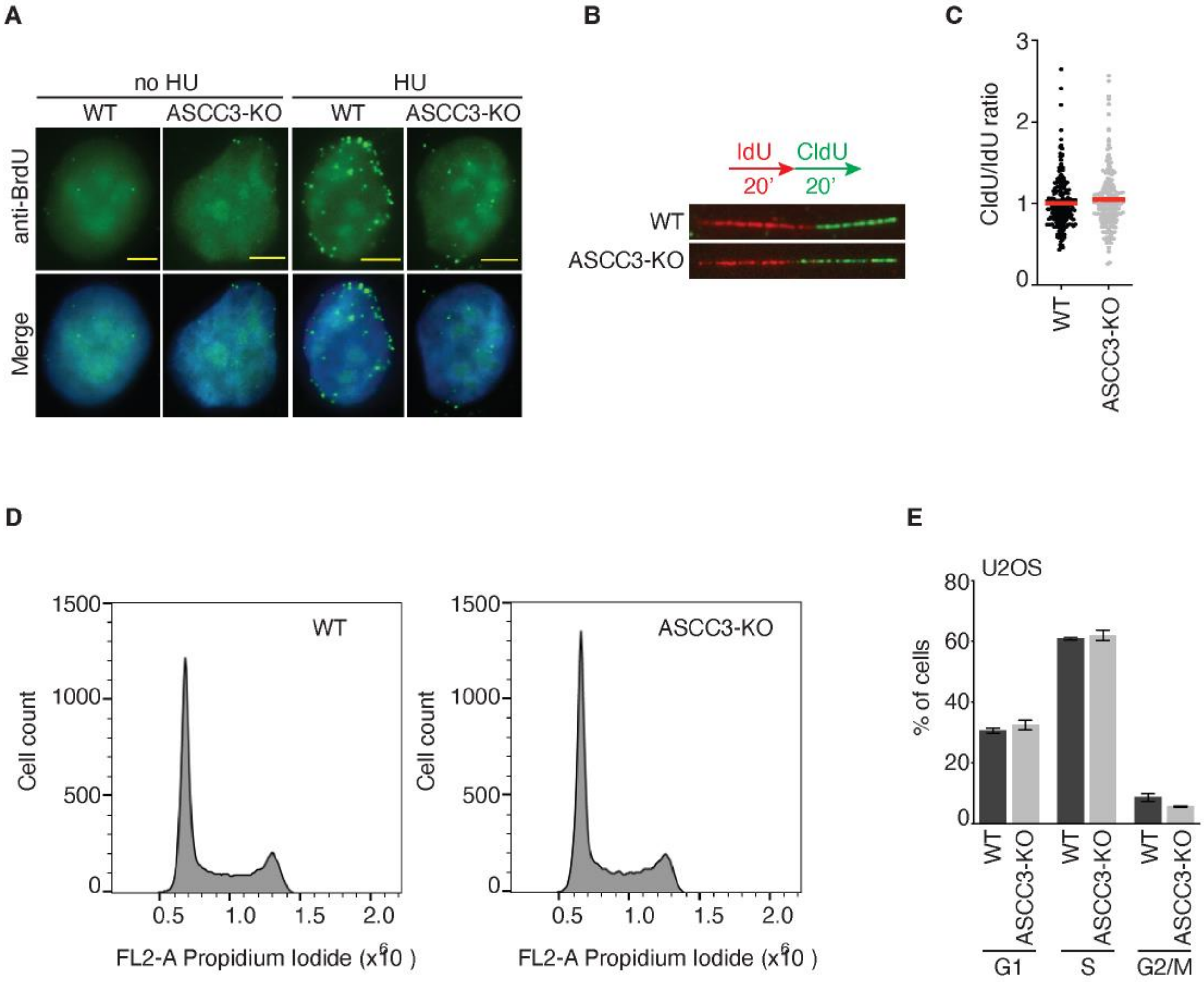

**Figure S4**. ASCC3 unwinds DNA upon replication stress. (**A**) Representative images of native BrdU staining in U2OS WT and ASCC3-KO cells that were pulse-labeled with BrdU prior to treatment with or without HU. (**B**) Representative images of DNA fibers from U2OS WT or ASCC3-KO cells that were first labeled with IdU and then CldU. (**C**) Quantification of the CldU/IdU ratio in U2OS WT and ASCC3-KO cells. This DNA fiber experiment was performed once. Data are represented as scatter plot graphs with the mean indicated. A total of 200-209 fibers per condition were analyzed. (**D**) Representative cell cycle profiles of U2OS WT and ASCC3-KO cells. (**E**) Quantification of the percentage of U2OS WT and ASCC3-KO cells in G1, S, and G2/M phases. SDs from three independent experiments are indicated.

Supplementary Figure S5 Cui et al

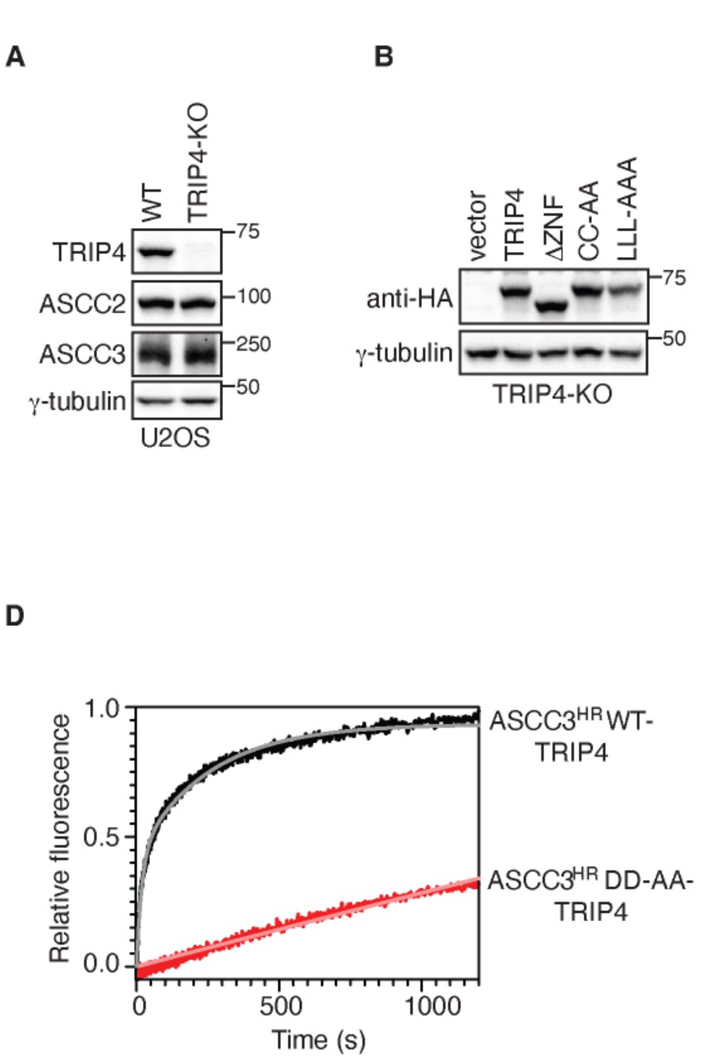

**Figure S5**. ASCC3 requires TRIP4 to promote RPA accumulation on ssDNA upon replication stress. (**A**) Western blot analyses of U2OS WT and TRIP4-KO cells. Immunoblotting was performed with anti-TRIP4, anti-ASCC2, anti-ASCC3, and anti-γ-tubulin antibodies. (**B**) Western blot analysis of U2OS TRIP4-KO cells expressing various HA-TRIP4 alleles as indicated. Immunoblotting was performed with anti-HA and anti-γ-tubulin antibodies. (**C**) Quantification of the percentage of EdU+ cells with ≥ 30 RPA32 foci in U2OS TRIP4-KO cells expressing various HA-TRIP4 alleles as indicated. These cells were pulse-labeled with EdU for 10 min prior to their treatment with or without HU. A total of 898-989 cells per condition were scored in blind. SDs from three independent experiments are shown. $**P<0.01$. (**D**) In vitro helicase assays of $ASCC3^{HR}$ WT-TRIP4 or $ASCC3^{HR}$-DD-AA-TRIP4 as indicated.

**Supplementary Figure S6 Cui et al**

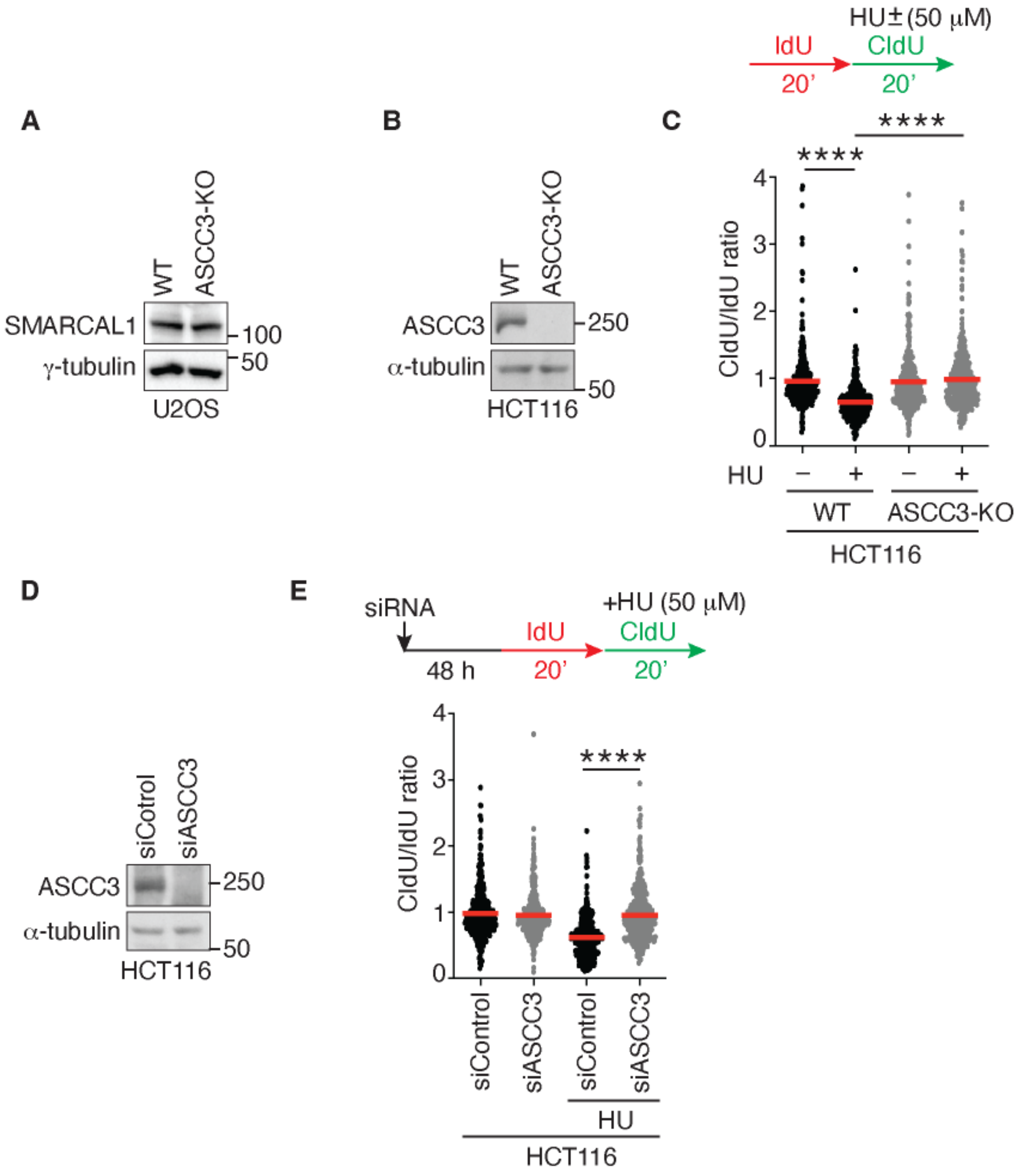

**Figure S6**. ASCC3 restrains fork progression upon replication stress. (**A**) Western blot analyses of U2OS WT and ASCC3-KO cells. Immunoblotting was performed with anti-SMARCAL1 and anti-γ-tubulin antibodies. (**B**) Western blot analyses of HCT116 WT and ASCC3-KO cells. Immunoblotting was performed with anti-ASCC3 and anti-α-tubulin antibodies. (**C**) Quantification of the CldU/IdU ratio in HCT116 WT and ASCC3-KO cells treated with or without HU. The DNA fiber experiments were performed twice independently with reproducible data in this and S6E panels. Data from one representative experiment are shown as scatter plot graphs with the mean indicated in this and S6E panels. A total of 308-317 fibers per condition were analyzed. The *P*-value was determined using a non-parametric Mann-Whitney rank-sum *t*-test in this and subsequent panels. ****$P<0.0001$. (**D**) Western blot analyses of HCT116 transfected with siControl or siASCC3. Immunoblotting was performed with anti-ASCC3 and anti-α-tubulin antibodies. (**E**) Quantification of the CldU/IdU ratio in HCT116 cells that were transfected with siControl or siASCC3 prior to treatment with or without HU. A total of 407-413 fibers per condition were analyzed. ****$P<0.0001$.

**Supplementary Figure S7 Cui et al.**

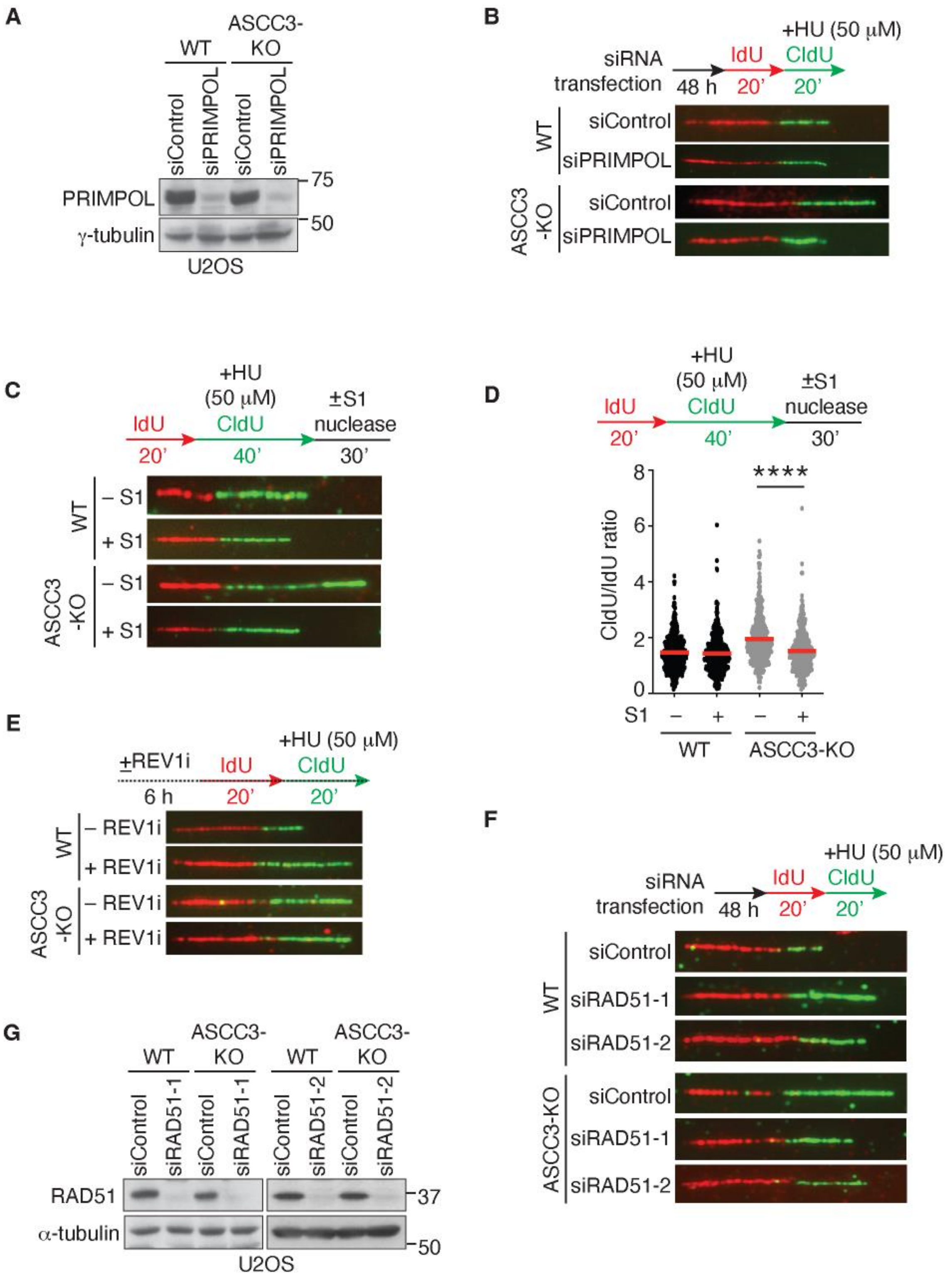

**Figure S7**. ASCC3 inhibits PRIMPOL- and RAD51-mediated damage tolerance pathways to restrain fork progression upon replication stress. (**A**) Western blot analyses of U2OS WT and ASCC3-KO cells transfected with control siRNA or siRNA against PRIMPOL (siPRIMPOL). Immunoblotting was performed with anti-PRIMPOL and anti-γ-tubulin antibodies. (**B**) Representative images of DNA fibers from HU-treated U2OS WT or ASCC3-KO cells that were transfected with control siRNA or siRNA against PRIMPOL. (**C**) Representative images of DNA fibers from HU-treated U2OS WT or ASCC3-KO cells that were treated with or without S1 nuclease. (**D**) Quantification of the CldU/IdU ratio in U2OS WT and ASCC3-KO cells. Following HU treatment, these cells were treated with or without S1 nuclease. The DNA fiber assays were performed twice independently with reproducible data. Data from one representative experiment are shown as scatter plot graphs with the mean indicated in this and subsequent panels. A total of 410-414 fibers per condition were analyzed. The *P*-value was determined using a non-parametric Mann-Whitney rank-sum *t*-test. ****$P$<0.0001. (**E**) Representative images of DNA fibers from HU-treated U2OS WT or ASCC3-KO cells that were treated with or without REV1 inhibitor (REV1i). (**F**) Representative images of DNA fibers from HU-treated U2OS WT or ASCC3-KO cells that were transfected with control siRNA or two independent siRNA against RAD51. (**G**) Western blot analyses of U2OS WT and ASCC3-KO cells transfected with indicated siRNAs. Immunoblotting was performed with anti-RAD51 and anti-γ-tubulin antibodies.

**Supplementary Figure S8 Cui et al.**

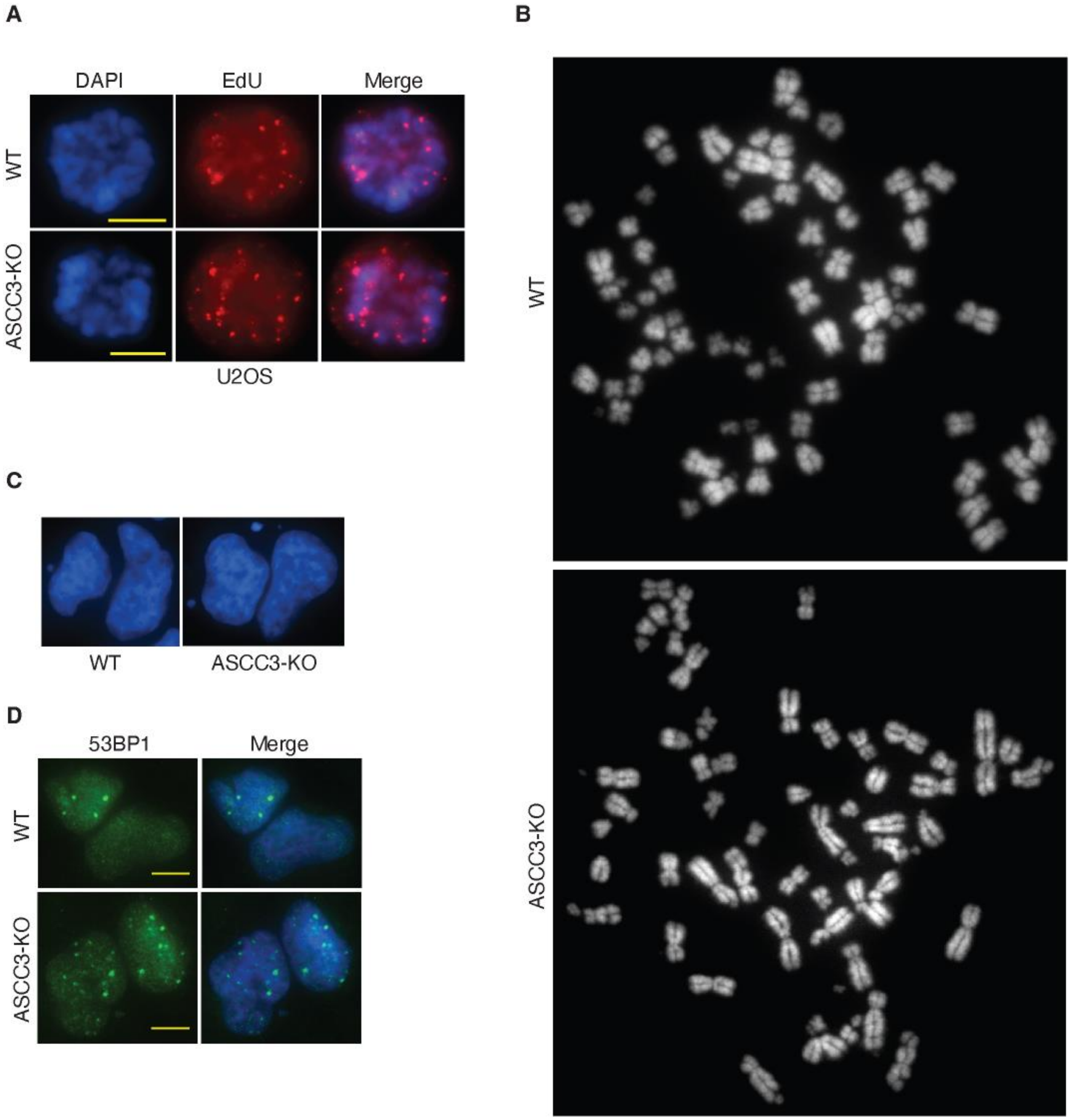

**Figure S8**. ASCC3 promotes genomic stability. (**A**) Representative images of EdU foci in U2OS WT and ASCC3-KO prometaphase cells following exposure to 0.4 μM aphidicolin in S phase. (**B**) Representative images of metaphase chromosomes of U2OS WT and ASCC3-KO cells following exposure to 0.4 μM aphidicolin in S phase. (**C**) Representative images of micronuclei formation in U2OS WT and ASCC3-KO G1 daughter cells following exposure to 0.4 μM aphidicolin in S phase. (**D**) Representative images of the formation of 53BP1 nuclear bodies in U2OS WT and ASCC3-KO G1 daughter cells following exposure to 0.4 μM aphidicolin in S phase.

Supplementary Figure S9 Cui et al.

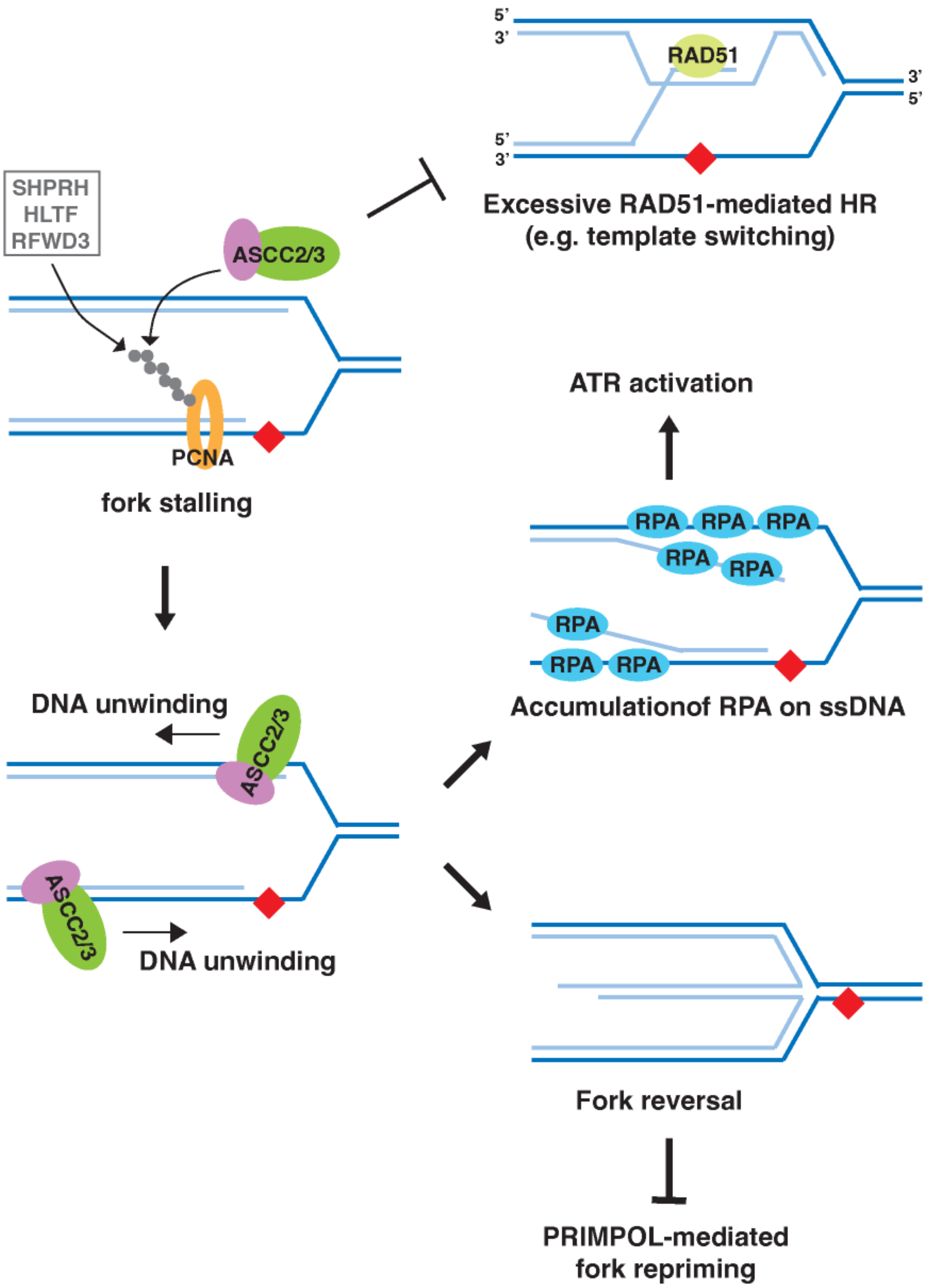

**Figure S9**. Model for the role of ASCC3 in controlling replication stress responses. See the text for details.
